# The antipsychotic drug clozapine suppresses autoimmunity driving psychosis-like behavior in mice

**DOI:** 10.64898/2026.03.28.714971

**Authors:** Le He, Harriet Feldman, Timothy Nguyen, Marion Bosc, Vasishta Polisetty, Orla Kriel, Antonia Landwehr, Annabel Borg, Fernanda Teixera Subtil, Mohammadparsa Khakpour, Jitong Zhou, Svend Kjær, James MacCabe, Thomas A. Pollak, Marie-Eve Tremblay, Carola G. Vinuesa, Adrian Hayday, Katharina Schmack

## Abstract

Antipsychotic drugs are the first-line treatment for psychosis yet their mechanism of action remains poorly understood, largely due to the challenge to faithfully model psychosis preclinically. Here, we focus on the emerging concept that psychosis can be caused by brain autoimmunity and present a novel mouse model of anti-N-methyl-D-aspartate-receptor (anti-NMDAR) encephalitis, a condition that manifests with psychosis and autoantibodies against the NMDAR. We devised a new mRNA-based approach to immunize mice against the NMDAR. Immunized mice developed psychosis-like behaviors that were caused by anti-NMDAR autoantibodies leading to phagocytosis of NMDARs by brain microglia. The antipsychotic drug clozapine rescued psychosis-like behaviors and, remarkably, reduced anti-NMDAR autoantibody levels and antibody-mediated phagocytosis of NMDARs. The immunomodulatory effects of clozapine were confirmed in a mouse model of systemic lupus erythematosus. Our results demonstrate that clozapine suppresses autoimmunity driving psychosis-like behaviors, raising the possibility that immunomodulation contributes to antipsychotic drug action.

**HIGHLIGHTS:**

- mRNA immunization against the NMDAR induces psychosis-like behavior in mice
- Anti-NMDAR autoantibodies are sufficient for psychosis-like behavior
- Microglial phagocytosis of NMDARs mediates psychosis-like behavior induced by anti-NMDAR autoantibodies.
- Clozapine reduces anti-NMDAR autoantibodies, microglial phagocytosis and psychosis-like behavior, consistent with immunomodulation as a potential mechanism of antipsychotic drug action.

## INTRODUCTION

> *“The schizophrenic mind is not so much split as shattered. I like to say schizophrenia is like a waking nightmare.” - Elyn Saks*

Schizophrenia is characterized by psychosis – a profound change in how the brain constructs reality manifesting with hallucinations, delusions, agitation and disorganized behavior. Psychotic disorders such as schizophrenia affect more than 20 million people worldwide, and continue to cause substantial global disease burden.^1^ This is notwithstanding the fact that first-line treatments – antipsychotic drugs – are effective at reducing the symptoms of acute psychosis,^2^ and can lead to lasting remission in some individuals. However, the overall benefits of anti-psychotic drugs are limited by significant adverse effects and high discontinuation rates, which contribute to poor long-term outcomes often observed in psychotic disorders.^3^ To advance treatments for psychotic disorders, it is crucial to distil the mechanisms through which antipsychotic drugs exert their benefits.

Despite decades of research, the mechanism of action of antipsychotic drugs remains incompletely understood. One crucial barrier to mechanistic investigations of antipsychotic drug action is the difficulty of recapitulating human psychosis in animal models amenable to experimental research. Because the exact biological changes that underpin acute psychosis symptoms remain elusive, most preclinical research has focused on modeling factors that increase the risk of developing psychosis – such as introducing human genetic variants associated with schizophrenia into mice.^4–6^ While such models have been invaluable for understanding disease susceptibility and synaptic biology, they are less suited to modeling the acute state of psychosis itself, which is in turn what antipsychotic drugs target. Here, we reasoned that the challenge of modeling acute psychosis in mice might be addressed by focusing on a rare autoimmune disease that presents with psychosis, namely anti-NMDAR encephalitis.

Using anti-NMDAR encephalitis as a model for acute psychosis offers the advantage of a well-defined, inducible biological cause that can be recapitulated in mice – autoantibodies targeting the glutamate NMDAR in the brain.^7^ The resulting NMDAR hypofunction is an established mechanism for psychosis: NMDAR antagonists induce psychosis-like states in humans^8^ and rare protein-truncating variants of the NMDAR substantially increase the risk for schizophrenia.^9^ In line with this, individuals with anti-NMDAR encephalitis present with acute psychosis in over 80% of the cases,^10,11^ before developing other overt neurological signs.^12,13^ Hence, mouse models of anti-NMDAR autoimmunity afford unique opportunities for a mechanistic dissection of acute psychosis and antipsychotic drug action.

Here, we present a new mouse model of anti-NMDAR autoimmunity to investigate the mechanisms by which antipsychotic drugs may treat acute psychosis. We introduce an mRNA-based approach to induce immune responses against the NMDAR and show that immunized mice develop behavioral alterations reminiscent of acute psychosis. Mechanistically, we demonstrate that this acute psychosis-like state is driven by autoantibodies binding to NMDARs, alongside microglial internalization of autoantibody–receptor complexes and reduced brain NMDAR levels. Using this model, we demonstrate that clozapine, the most effective antipsychotic drug,^2^ prevents the emergence of psychosis-like behaviors, and, strikingly, also reduces anti-NMDAR autoantibody levels. We confirm this immunomodulatory action of clozapine in a mouse model of lupus-like systemic autoimmunity, where clozapine similarly reduces autoantibody levels and ameliorates disease phenotypes. Together, these findings further establish anti-NMDAR autoimmunity as a mechanistic model of acute psychosis and suggest that clozapine may exert antipsychotic effects, at least in part, through immune modulation.

## RESULTS

### mRNA immunization against the NMDAR induces psychosis-like behavior in mice

We first sought to establish an experimental approach to induce anti-NMDAR autoimmunity in mice capable of recapitulating behavioral phenotypes of acute psychosis. To elicit a durable and comprehensive immune response against NMDAR, a large self-protein composed of multiple subunits, we used mRNA encapsulated in lipid nano-particles (LNP), drawing on recent advances in vaccine technology.^14^ We injected mice with mRNA-LNP encoding the three major subunits of the NMDAR (GluN1, GluN2A, GluN2B), or with mRNA-LNP encoding an exogenous control antigen (luciferase, **Fig. 1A**).

**Figure 1:**
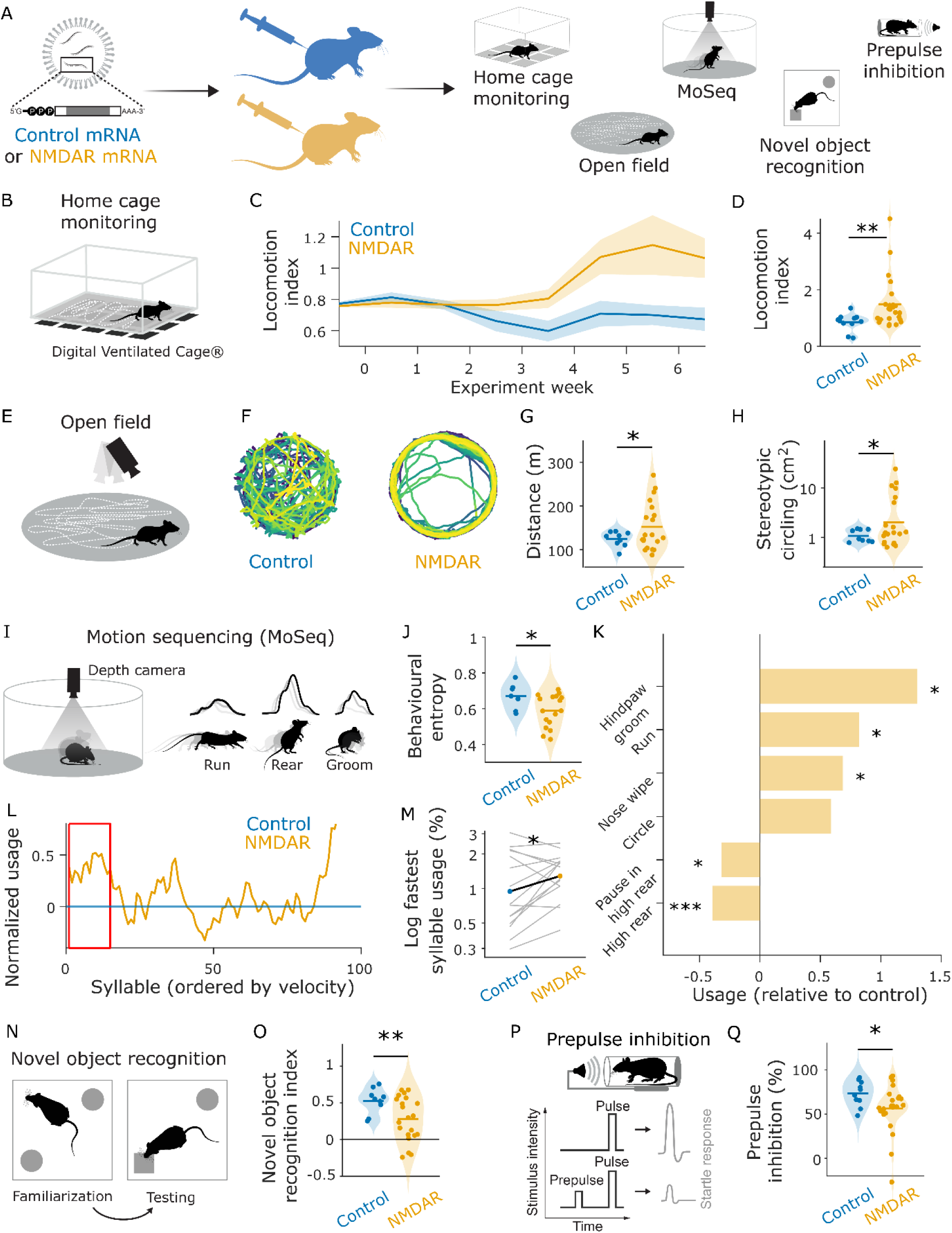
mRNA-LNP anti-NMDAR immunization induces psychosis-like behavior. (A) Experimental design. mRNA coding for NMDAR or luciferase (control) encapsulated in lipid nanoparticles was used. Mice received three injections spaced by two weeks. Behavior was assessed throughout. (B) Home cage monitoring setup (C) Home cage locomotion over time. (D) Home cage locomotion at endpoint was increased in NMDAR-immunized compared to control mice. (E) Open field experimental setup. (F) Representative trajectories of two exemplary mice at endpoint. (G) Distance travelled was increased in NMDAR-immunized compared to control mice. (H) Stereotypical circling was increased in NMDAR immunized compared to control mice (I) Motion Sequencing (MoSeq) experimental setup. (J) Behavioral entropy, measured as syllable transition matrix entropy, was reduced in NMDAR-immunized compared to control mice, indicating narrowed behavioral repertoire. (K) Usage of running and grooming syllables was increased and rearing decreased, in NMDAR-immunized mice normalized to control mice. Bars: mean. Overall linear mixed effects model of group x category; interaction p<0.001. Bars: mean. Stars: post-hoc t-test for each category. (L) Usage of syllables in NMDAR-immunized mice relative to control mice. Red square denotes the fastest syllables analyzed in (M) (M) Usage of fastest syllable was increased in NMDAR-immunized mice compared to control mice, consistent with findings from a Grin2a knockout model. Dots: mean across syllables. Grey lines: syllables. Statistical test: paired t-test. (N) Novel object recognition experimental setup. (O) Novel object recognition index was reduced in NMDAR-immunized mice compared to control mice, indicating reduced preference for novel objects. (P) Prepulse inhibition experimental setup. (Q) Prepulse inhibition, calculated as the attenuation of the acoustic startle response on prepulse compared to no prepulse trials, was reduced in NMDAR-immunized mice compared to control mice. All dot plots: dots represent individual animals; horizontal line indicates group mean; shaded area indicates probability density. Time-series panels: lines indicate group means; shaded areas indicate SEM. Group comparisons by Welch’s t-test throughout unless otherwise noted. ∗p < 0.05, ∗∗p < 0.01, ∗∗∗p < 0.001.

We noticed that mice immunized against the NMDAR started to display unusual behavior that was clearly apparent upon visual inspection (**Supplementary Videos 1-4**). Building on these qualitative observations, we proceeded to using digital ventilated cages to continuously record home cage activity (**Fig. 1B)**. We found that NMDAR-immunized mice consistently developed hyperlocomotion (**Fig. 1C-D**), mirroring the agitation often seen in acute psychosis. These findings were corroborated in the open field (**Fig. 1E**), where NMDAR-immunized mice but not controls again showed hyperlocomotion alongside stereotypical circling behavior (**Fig. 1F-H**). These results confirmed that immunization against the NMDAR elicited a behavioral phenotype recapitulating key psychomotor features of acute psychosis.

To further characterize the behavioral phenotype elicited by immunization against the NMDAR, we used an unsupervised machine learning approach which parses video recordings of behavior into sub-second ‘syllables’ (MoSeq^15^, **Fig. 1I**). Here, analyses of behavioral sequences revealed lower entropy in NMDAR-immunized compared to controls (**Fig. 1J**), reflecting a more constrained behavioral repertoire. NMDAR-immunized mice relative to controls showed decreases in rearing and increases in grooming, running and circling (**Fig. 1K**), indicating a contraction of exploratory behaviors in favor of stereotypical behaviors. Interestingly, we observed an increase in the use of the fastest syllables (**Fig. 1L-M**), similar to what has been reported in a genetic mouse model of schizophrenia with a knockout of an NMDAR subunit,^6,16^ lending independent validation for our mouse model of psychosis grounded in anti-NMDAR autoimmunity.

To situate our work within the broader literature on anti-NMDAR autoimmunity and psychosis, we next employed established behavioral assays of cognitive and sensorimotor function. Replicating previous reports,^17–19^ NMDAR-immunized mice relative to controls exhibited less preference for novel over familiar objects (**Fig. 1N-O**), consistent with memory deficits found both in anti-NMDAR encephalitis^13^ and schizophrenia.^20^ Furthermore, NMDAR-immunized mice compared to controls showed reduced pre-pulse inhibition (**Fig. 1P-Q**), a cross-species biomarker of sensorimotor gating consistently reduced in patients with schizophrenia.^21^

Collectively, these findings demonstrate that mRNA immunization against the NMDAR induces robust behavioral phenotypes in mice capturing several aspects of acute psychosis.

### Anti-NMDAR autoantibodies induce psychosis-like behavior

We next sought to understand the mechanisms by which immunization against the NMDAR gives rise to the observed psychosis-like behavior. Anti-NMDAR encephalitis is considered an antibody-mediated disease,^7^ consistent with which, NMDAR-immunized mice but not controls developed serum anti-NMDAR autoantibodies (**Fig. 2A-B**). Serum levels of anti-NMDAR autoantibodies correlated positively with locomotion across animals (**Fig. 2C**) and tracked locomotion over time (**Fig. 2D**). More broadly, anti-NMDAR autoantibody levels correlated in the expected direction with the majority of the behavioral phenotypes reported above (**Fig. 2E**), indicating a robust relationship between anti-NMDAR autoantibodies and psychosis-like behavior.

**Figure 2:**
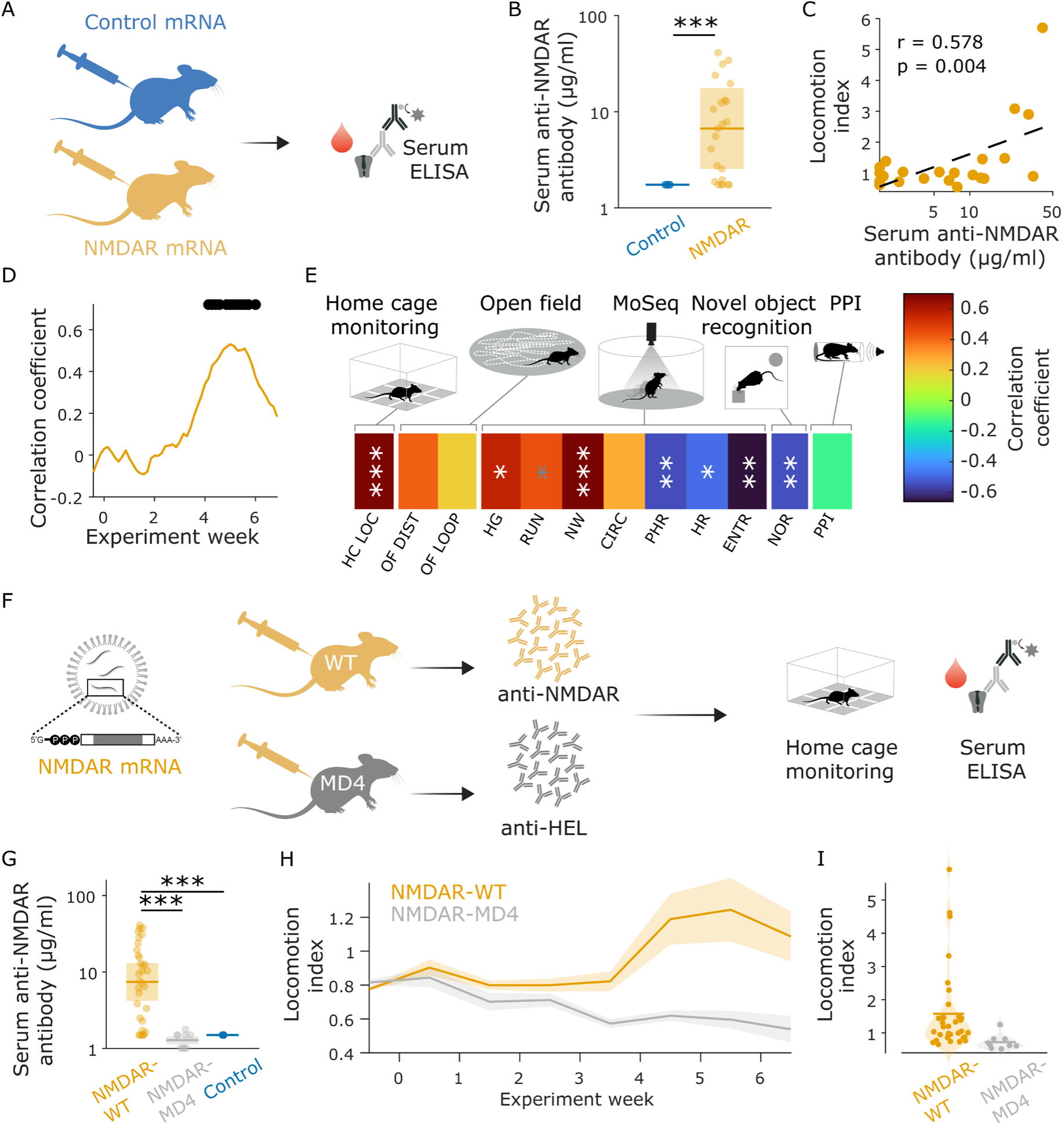
Anti-NMDAR antibodies are related to psychosis-like behavior. (A) Experimental design for antibody measures. NMDAR- or luciferase(control)-encoding mRNA-LNP was administered in two injections two weeks apart; serum antibody levels were measured one week later with by a custom ELISA. (B) Anti-NMDAR autoantibody serum concentrations were elevated in NMDAR-immunized compared to control mice. Dots represent individual animals. Shaded area: interquartile range. Horizontal bar: median. Group comparison by Wilcoxon’s rank sum test. (C) Correlation between log anti-NMDAR autoantibody concentration with home cage locomotion at maximum timepoint (day 37). Dots: individual animals. Dashed line: least-square regression. Statistical test: Pearson correlation. (D) Time course of correlation between log anti-NMDAR autoantibody concentration and home cage locomotion. Black dots indicate days where p<0.05. (E) Correlation matrix between log anti-NMDAR autoantibody concentrations and all behavioral metrics shown in Figure 1. HC LOC – Homecage locomotion, OF DIST – Open field distance, OF LOOP – Open field stereotypical circling, HG – MoSeq hindpaw groom, RUN – MoSeq run, NW – MoSeq nose wipe, CIRC – MoSeq circling, PHR – MoSeq pause in high rear, HR – MoSeq high rear, ENTR – MoSeq behavioral entropy, NOR – novel object recognition index, PPI – prepulse inhibition. Statistical test: Pearson correlation. Grey * p<0.1. ∗p < 0.05, ∗∗p < 0.01, ∗∗∗p < 0.001. (F) Experimental design to probe antibody necessity. Wild-type (WT) and MD4 mice, which constitutively produce antibodies against hen egg lysozyme (HEL) and therefore have a limited capacity to mount specific antibody responses to other antigens, were immunized against NMDAR in three injections spaced two weeks apart. Home cage behavior was monitored throughout; serum antibody levels were assessed by custom ELISA at endpoint (G) MD4 mice compared to WT mice showed lower anti-NMDAR autoantibody concentration, comparable to controls. Dots represent individual animals. Shaded area: interquartile range. Horizontal bar: median. Group comparisons by Wilcoxon’s rank sum test. (H) Home cage locomotion over time in MD4 and WT mice. (I) MD4 mice compared to WT mice showed decreased home cage locomotion at endpoint. Dots represent individual animals; horizontal line indicates group mean; shaded area indicates probability density. Group comparisons by Welch’s t-test. All dot plots: dots represent individual animals; horizontal line indicates group mean; shaded area indicates ±1SD (B and G) or probability density (I). Time-series panels (H): lines indicate group means; shaded areas indicate SEM.∗p < 0.05, ∗∗p < 0.01, ∗∗∗p < 0.001.

We next tested whether anti-NMDAR autoantibodies were *necessary* for psychosis-like behavior. To this end, we employed MD4 transgenic mice which when immunized against an antigen such as the NMDAR, are unable to generate a specific antibody response because they carry a fixed B cell receptor for hen egg lysozyme (**Fig. 2F**). In line with this, NMDAR-immunized MD4 mice in contrast to wild-type mice did not develop any anti-NMDAR auto-antibodies (**Fig. 2G**). Critically, NMDAR-immunized MD4 mice did not show the hyperlocomotion observed in NMDAR-immunized WT mice (**Fig. 2H-I**), indicating a crucial role for NMDAR-specific B cells. While antigen-specific B cells have several functions in addition to antibody production, these findings are consistent with anti-NMDAR antibodies being necessary for psychosis-like behavior.

We next asked if anti-NMDAR autoantibodies, once formed, might be *sufficient* to induce psychosis-like behavior. Thus, we passively transferred serum IgG harvested from NMDAR-immunized mice (anti-NMDAR IgG) or control-immunized mice (control IgG) to naïve recipient mice through constant infusion into the lateral ventricles via implanted osmotic minipumps (**Fig. 3A**). Anti-NMDAR IgG recipients but not control IgG recipients developed detectable serum levels of anti-NMDAR autoantibodies (**Fig. 3B**), confirming that passive transfer had occurred.

**Figure 3:**
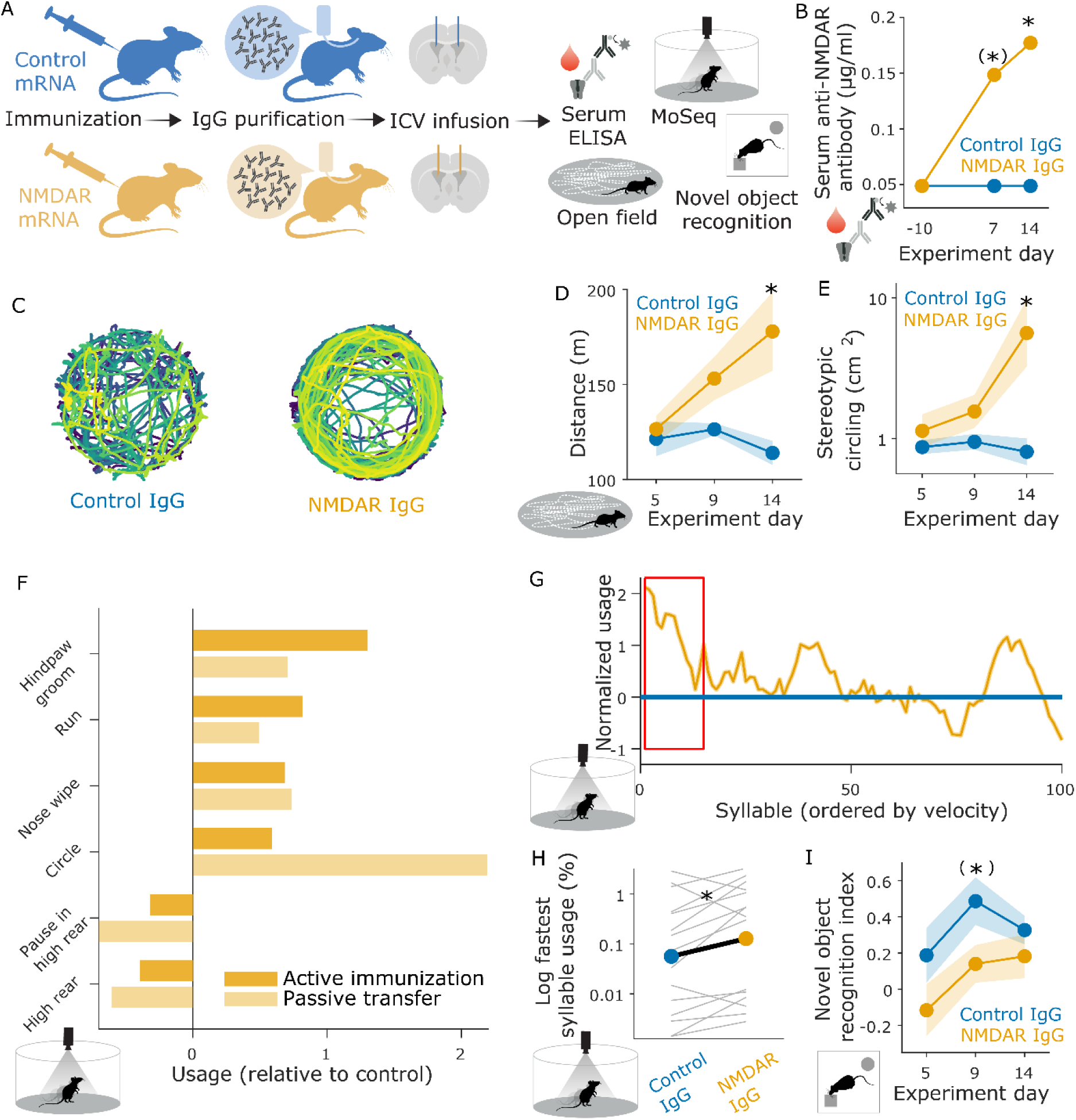
Anti-NMDAR antibodies are sufficient for psychosis-like behavior. (A) Passive transfer experimental design. IgG was purified from serum of NMDAR- or luciferase (control)-immunized donor mice and delivered continuously to naive recipient mice via intracerebroventricular (ICV) osmotic minipumps over two weeks. Behavior and serum antibody levels were assessed throughout. (B) Anti-NMDAR autoantibody concentrations in serum increased over time in NMDAR IgG recipients but not in control IgG. Dots: group median. (∗)p < 0.1, ∗p < 0.05, Wilcoxon rank sum test. (C) Representative open field trajectories of two mice on day 14 after minipump implantation. (D) Open field locomotion was increased in NMDAR IgG recipients compared to control IgG recipients. Overall linear mixed effects model of group x timepoint interaction p < 0.05. ∗p < 0.05 in post-hoc t-test. Dots: individual animals. Shaded areas: Cosineau-corrected SEM. (E) Open field stereotypic circling was increased in NMDAR IgG recipients compared to control IgG recipients. Overall linear mixed effects model of group x timepoint interaction p < 0.05. ∗p < 0.05 in post-hoc t-test. Data types as in (D) (F) MoSeq behavioral syllable usage was altered in NMDAR IgG recipients relative to control IgG recipients, in a pattern consistent with the mRNA active immunization experiment. Overall linear mixed effects model of group x category p <0.05 (passive transfer experiment only). (G) and (H) Greater use of fastest syllable in NMDAR IgG recipients compared to control IgG recipients, in line with results from a Grin2a knockout model^5,16^ and mRNA immunization experiment. (H) Dots: mean across syllables. Grey lines: syllables. Statistical test: paired t-test. (I) NMDAR IgG recipients compared to control IgG recipients show a transient trend towards decreased preference for novel objects 9 days after implantation. Overall linear mixed effects model of group and timepoint with main effect group p <0.05 and main effect timepoint p<0.05. (∗)p <0.1 in post-hoc t-test. (∗) p<0.1 ∗p < 0.05, ∗∗p < 0.01, ∗∗∗p < 0.001.

Strikingly, passive transfer of IgG from NMDAR-immunized mice reproduced most of the behavioral phenotypes observed after active immunization against NMDAR. While home cage activity was not meaningfully interpretable due to large individual variability in implant-related movement restriction, anti-NMDAR IgG recipients but not control IgG recipients developed hyperlocomotion and stereotypical circling in the open field (**Fig. 3C-E**), closely mirroring NMDAR-immunized mice (cf. Fig 1H-J). Moreover, behavioral syllable analyses using MoSeq revealed that anti-NMDAR IgG recipients relative to control IgG recipients showed a shift in their behavioral repertoire similar to that seen in NMDAR-immunized mice: decreases in exploratory syllables – rearing – and increases in stereotypical syllables – grooming, running and circling (**Fig. 3F**, cf. Fig 1M), which was paralleled by an increased use of high-speed syllables (**Fig. 3G-H**, cf. Fig 1 N-O). Likewise, novel object recognition was overall decreased in anti-NMDAR IgG recipients compared to control IgG recipients, consistent with our findings in NMDAR-immunized mice (**Fig. 3I**, cf. Fig. 1Q). Pre-pulse inhibition could not be assessed in this cohort owing to the risk of minipump displacement.

In sum, these results indicate that anti-NMDAR autoantibodies are necessary and sufficient to induce psychosis-like behavior in mice.

### Anti-NMDAR autoimmunity manifests with CNS autoantibody deposition and NMDAR loss

To understand how anti-NMDAR autoantibodies might cause psychosis-like behavior, we first asked whether anti-NMDAR autoantibodies gain access to the brain to directly interfere with their antigenic target, the NMDAR. In cerebrospinal fluid, anti-NMDAR autoantibodies were detected in NMDAR-immunized mice but not controls (**Fig. 4A-B**), indicating that autoantibodies were present within the CNS compartment. Moreover, cerebrospinal fluid anti-NMDAR autoantibody levels correlated with home cage locomotion (**Fig. 4C**). Immunofluorescence microscopy of brain tissue revealed IgG deposition and reduced NMDAR levels in anti-NMDAR mice relative to control mice (**Fig. 4D-F**), and the results were confirmed by Western blot (**Fig. 4G-I**). These findings are consistent with anti-NMDAR autoantibodies gaining access to the CNS and causing loss of NMDARs.

**Figure 4:**
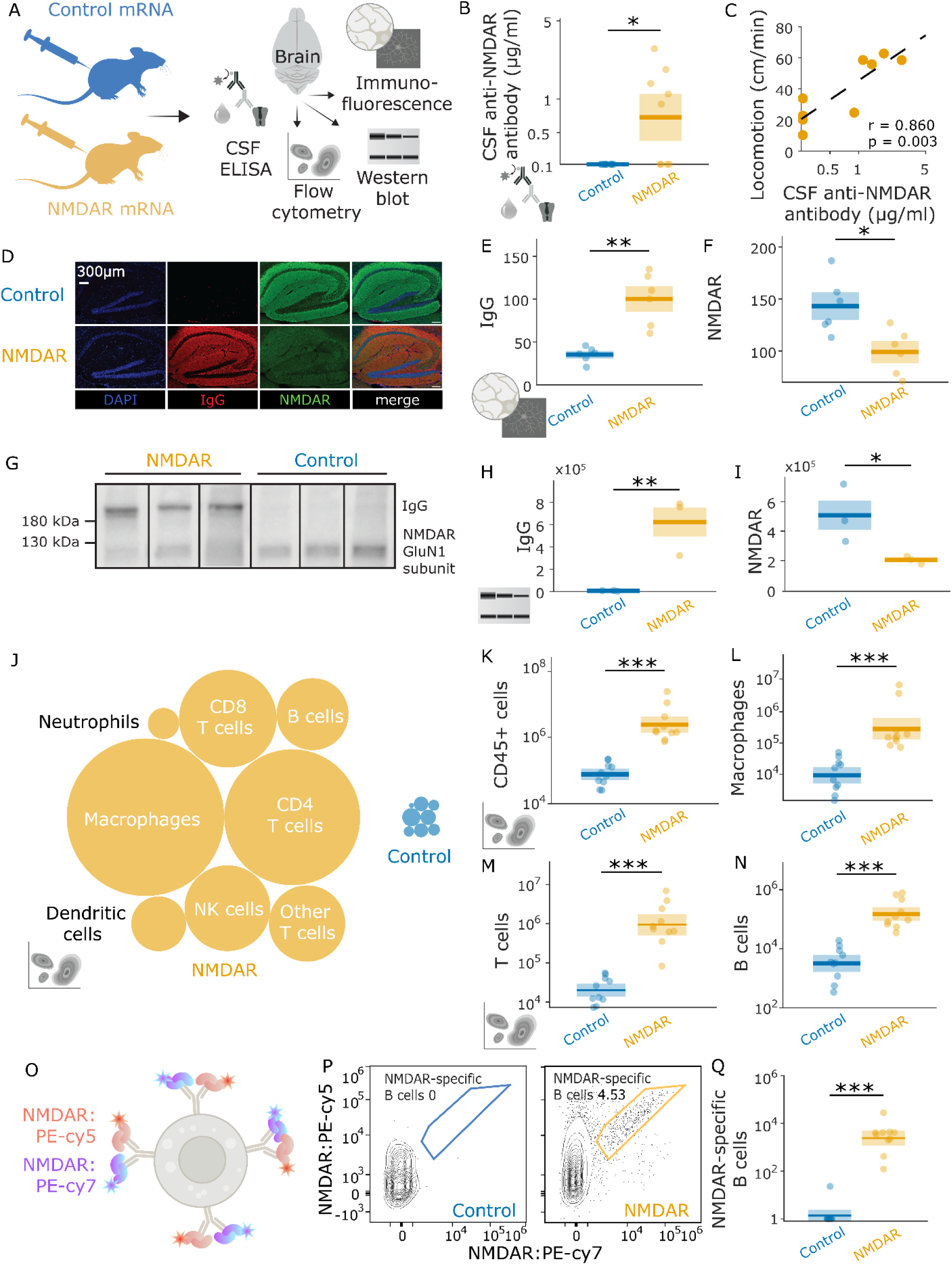
Anti-NMDAR autoimmunity manifests in the brain with CNS antibody production and NMDAR loss. (A) Experimental design. NMDAR- or control mRNA was administered in two injections two weeks apart; brains were collected one week later and assessed with immunofluorescence microscopy image analysis, Western blot and flow cytometry. (B) Anti-NMDAR autoantibody in CSF was higher in NMDAR-immunized mice compared to control mice. Dots represent individual animals. Shaded area: interquartile range. Horizontal bar: median. Group comparison by Wilcoxon’s rank sum test. (C) CSF anti-NMDAR autoantibody concentration correlated with home cage locomotion at endpoint. Dots: individual animals. Dashed line: least-square regression. Statistical test: Pearson correlation. (D) Representative sagittal brain sections from one NMDAR-immunized and one control mouse, stained for DAPI, IgG and NMDAR. NMDAR immunofluorescence is distributed across the neuropil, reflecting the synaptic localization of the receptor, and is spatially distinct from DAPI-stained nuclei. In NMDAR-immunized mice, IgG immunofluorescence follows the distribution of NMDAR immunofluorescence, which in turn is markedly reduced, consistent with autoantibody-mediated receptor loss. (E-F) Anti-IgG immunofluorescence was increased and anti-NMDAR immunofluorescence decreased in the brains of NMDAR-immunized mice compared to control mice. (G) - (I) IgG levels were increased and NMDAR levels decreased in hippocampal Western blots of NMDAR-immunized mice compared to control mice. (J) Immune cell populations in the brain of NMDAR-immunized and control mice as revealed by flow cytometry. Circle size indicates relative population size. Live singlet CD45⁺ cells were gated sequentially: macrophages (CD45-Hi, F4/80⁺CD64⁺); dendritic cells (CD45-Hi, F4/80⁻CD64⁻, MHC-II⁺CD11c⁺); NK cells (NK1.1⁺Ly6G⁻); neutrophils (NK1.1⁻Ly6G⁺); B cells (CD19⁺CD3⁻); CD4⁺ T cells (CD3⁺CD4⁺CD8⁻); CD8⁺ T cells (CD3⁺CD4⁻CD8⁺); and other T cells (CD3⁺CD4⁻CD8⁻). (K)-(N) Immune cell counts in the brain of NMDAR-immunized mice compared to control mice. Immune cell populations were identified as described in (J). (O) Experimental approach for identification of NMDAR-specific B cells. The NMDAR antigen was conjugated to two spectrally distinct fluorophores (PE-Cy5 and PE-Cy7); B cells binding both were classified as NMDAR-specific B-cells. (P) Representative contour flow cytometry plots from one NMDAR-immunized and one control mouse, showing NMDAR-specific B cells. (Q) Antigen-specific B cells were present at higher numbers in the brains of NMDAR-immunized compared to control mice. All dot plots: unless otherwise specified, dots represent individual animals; horizontal line indicates group mean; shaded area indicates ± 1 SD. Group comparisons by Welch’s t-test throughout unless otherwise noted. ∗p < 0.05, ∗∗p < 0.01, ∗∗∗p < 0.001.

To better understand the immune processes leading to anti-NMDAR autoantibodies in the CNS, we next characterized immune cells in the brain with flow cytometry. NMDAR-immunized mice compared to controls exhibited higher numbers of immune cells in the brain (**Fig. 4J**), including innate cell lineages, T cells and B cells (**Fig. 4K-N**), raising the possibility that local antibody synthesis may contribute to CNS anti-NMDAR autoantibody deposition. To test this possibility, we designed a dual-labeling approach for measuring NMDA-receptor-specific B cells (**Fig. 4O**). This revealed the presence of NMDAR-specific B cells in the brain of NMDAR-immunized mice but not controls (**Fig. 4P-Q**), in line with a local cellular source of anti-NMDAR autoantibodies in the CNS.

In summary, these findings are consistent with a model in which autoantibodies provoke a loss of NMDARs in the brain, and the resulting NMDAR hypofunction leads to psychosis-like behavior, mirroring observations from genetic and pharmacological NMDAR disruption.^5,6,16,22^

### Microglial NMDAR phagocytosis contributes to psychosis-like behavior

We next asked how anti-NMDAR autoantibodies may lead to NMDAR loss. We focused on microglia, based on recent work showing microglial activation and NMDAR phagocytosis in models of NMDAR autoimmunity.^22,23^ Using fluorescence microscopy (**Fig. 5A**), we observed striking morphological differences in microglia of NMDAR-immunized mice compared to controls (**Fig. 5B**) with quantitative decreases in filament length (**Fig. 5C**), consistent with a shift towards a reactive microglial state. We further observed colocalization of NMDAR with microglial and lysosomal markers Iba1 and CD68 in anti-NMDAR mice (**Fig. 5D**), suggestive of microglial NMDAR phagocytosis.^24,25^ Indeed, we corroborated the location of NMDAR inside lysosomes within microglia using immunoperoxidase electron microscopy, finding microglial endosomes densely packed with NMDAR in NMDAR-immunized mice (**Fig. 5E**). Taken together, these findings confirm microglial NMDAR phagocytosis in anti-NMDAR autoimmunity.

**Figure 5:**
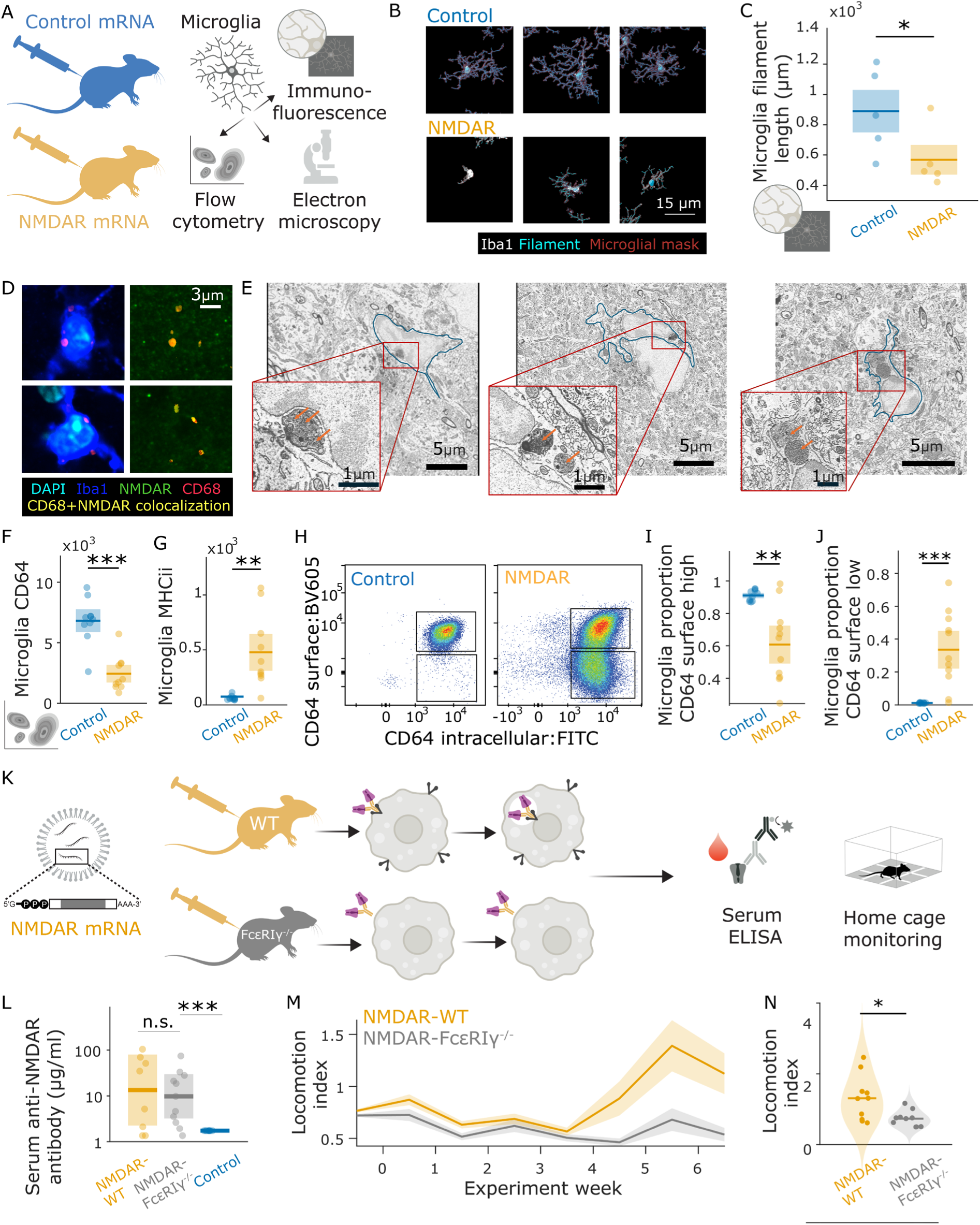
Antibody-mediated phagocytosis causes psychosis-like behavior in NMDAR-immunized mice. (A) Experimental design. NMDAR- or control mRNA was administered in two injections two weeks apart; brains were collected one week later and assessed with confocal immunofluorescence microscopy, immunoelectron microscopy and microglia-specific flow cytometry. (B) Representative confocal images stained for Iba1 to delineate microglia. (C) Microglia filament length, a marker of resting state, was decreased in NMDAR-immunized mice compared to control mice. Statistical significance was assessed in linear mixed effects model on cells with fixed factor group and random factor mouse. Dots: mean across cells per mouse. Horizontal lines: group means. Shaded areas: ± 1 group SD. (D) Representative confocal microscopy images showing accumulation of NMDAR within microglial lysosomes. Left panels: lysosomes (red) located around the nucleus of microglia. Right panels: NMDAR co-located with these lysosomes. Dark blue: IBA1 (microglia). Light blue: DAPI. Green: NMDAR. Red: CD68 (lysosome). Yellow: Colocalization of CD68 and NMDAR. (E) Representative immunoperoxidase electron microscopy images of hippocampal microglia from NMDAR-immunized mice, showing HRP-labelled NMDAR (orange arrows) concentrated in enlarged perinuclear lysosomes. Blue contours: microglial boundaries (F) - (G) Microglial expression of the IgG Fc gamma receptor FcγRI CD64 was reduced in NMDAR-immunized mice compared to control mice, while microglial expression of the activation marker MHC-II was increased. Microglia were identified within the CD45-Lo fraction as CD11b⁺. (H) Representative pseudocolour flow cytometry plots showing surface and intracellular CD64 staining in microglia, for one NMDAR-immunized mouse and one control mouse. Black boxes denote gates for quantification in (I) and (J) (I) – (J) Proportion of microglia with high intracellular CD64 shifted from a surface CD64-high (I) to a surface CD64-low (J) phenotype in NMDAR-immunized compared to control mice, suggesting CD64 internalization. (K) Experimental design. FcεRIγ-/-mice, which lack the FcR γ-chain required for surface expression and signaling of all activating Fcγ receptors and are therefore incapable of antibody-mediated phagocytosis, were immunized with NMDAR mRNA in three injections at week 0, 2 and 5. Home cage activity was monitored throughout. Serum anti-NMDAR antibody levels were assessed at endpoint. (L) Serum anti-NMDAR autoantibody concentrations were increased in both FcεRIγ^−/−^ and WT NMDAR-immunized mice compared to controls, confirming that the FcεRIγ knockout does not impair antibody production. Group comparison by Wilcoxon rank-sum test. (M) Home cage locomotion over time in FcεRIγ^−/−^ and WT NMDAR-immunized mice. (N) Home cage locomotion at endpoint was decreased in FcεRIγ^−/−^ compared to WT NMDAR-immunized mice. All dot plots: dots represent individual animals; horizontal line indicates group mean; shaded area indicates ±1SD (C, F, G, I, J) or probability density (N). Time-series panel (M): lines indicate group means; shaded areas indicate SEM. Group comparisons by Welch’s t-test throughout unless otherwise noted. ∗p < 0.05, ∗∗p < 0.01, ∗∗∗p < 0.001.

We next asked whether NMDAR phagocytosis is a causal mechanism by which anti-NMDAR autoantibodies drive psychosis-like behavior, rather than an epiphenomenal response to neuronal injury. We reasoned that if NMDAR phagocytosis is causally involved, disrupting the molecular machinery of antibody-dependent phagocytosis should attenuate psychosis-like behavior. Using flow cytometry, we measured microglial CD64, the major activating IgG Fc gamma receptor FcγRI that is an important mediator of antibody-dependent phagocytosis in myeloid cells including microglia.^23,24^ This revealed decreased CD64 on microglia in anti-NMDAR mice compared to controls (**Fig. 5F**), while the activation marker MHC-II, in contrast, was increased (**Fig. 5G**). The dissociation of these two molecular markers is consistent with reduced CD64 levels reflecting its internalization in the context of antibody-dependent phagocytosis. Indeed, staining intra- and extra-cellular CD64 revealed the emergence of a population of microglia with depleted surface CD64 but preserved internal CD64 after immunization (**Fig. 5H-J**). These results suggest that phagocytosis of NMDAR may be a key mechanism by which anti-NMDAR autoantibodies lead to psychosis-like behavior.

To test this hypothesis, we immunized mice which lack FcR gamma chain required for the surface expression and signaling of all activating FcγRs and which are therefore incapable of FcR-dependent, antibody-mediated phagocytosis (**Fig. 5K**). After immunization against NMDAR, FcεRIγ^−/−^ mice compared to WT mice mounted comparable levels of anti-NMDAR autoantibodies (**Fig. 5L**) yet showed attenuated psychosis-like behavior with significantly reduced hyperlocomotion (**Fig. 5M-N**). These findings implicate FcγR-mediated antibody-dependent phagocytosis as a causal contributor to psychosis-like behavior.

Collectively, our results establish that NMDAR phagocytosis contributes to the expression of psychosis-like behavior driven by anti-NMDAR autoantibodies.

### Clozapine reduces psychosis-like behavior in anti-NMDAR autoimmunity

Although anti-NMDAR encephalitis and schizophrenia are distinct conditions, they both present with psychosis and both involve NMDAR hypofunction. The mechanistic link we established between anti-NMDAR autoantibodies and psychosis-like behaviors therefore offers a well-characterized model for exploring antipsychotic drug action, starting with clozapine, the most effective antipsychotic drug. Mimicking clinical dosing practice, we gradually titrated mice onto a diet containing 750ppm clozapine before immunization against the NMDAR (**Fig. 6A**). Plasma clozapine levels in clozapine-treated mice (69 ± 11 ng/mL, data not shown) were in the order of plasma levels reported for patients taking clozapine,^27^ confirming that our dosing approach achieved clinically comparable clozapine exposure.

**Figure 6:**
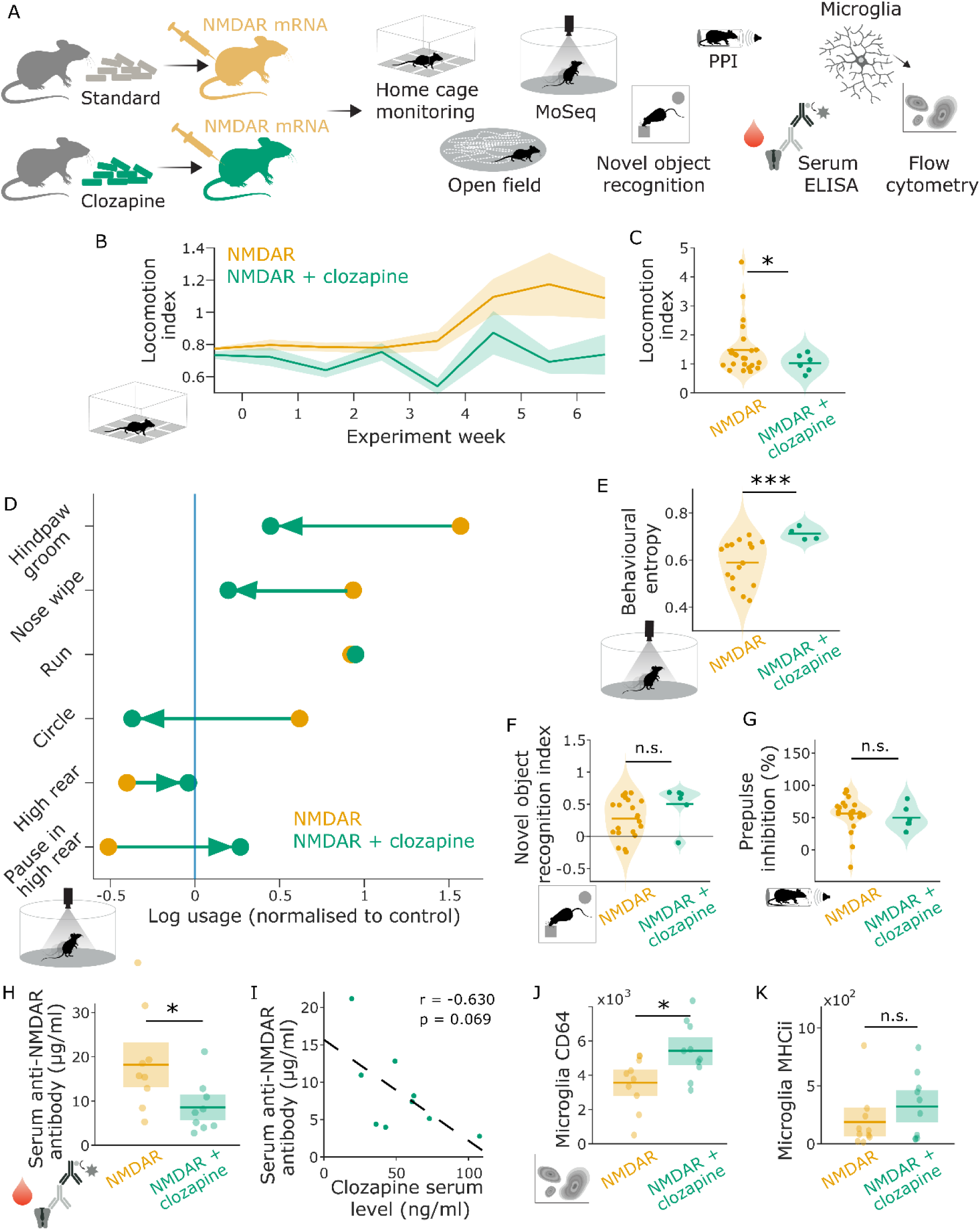
Clozapine reduces psychosis-like behavior and anti-NMDAR autoantibodies. (A) Experimental design. Mice were weaned on to a diet containing 750ppm clozapine or stayed on a standard diet before immunizations. NMDAR mRNA was administered in three injections two weeks apart while behavior was assessed in different assays, or in two injections two weeks apart before brains were collected for flow cytometry one week later. (B) Home cage locomotion over time in NMDAR-immunized mice fed clozapine or standard diet. (C) Home cage locomotion at endpoint was reduced in NMDAR-immunized mice fed clozapine compared to standard diet. (D) Clozapine diet partially normalized MoSeq syllable usage in NMDAR-immunized mice toward control levels. Yellow dots: NMDAR-immunized mice on control diet. Green dots: NMDAR-immunized mice on clozapine diet. Blue line: control-immunized mice on control diet (reference, normalized to 0). Linear mixed effects model of group x category; interaction p<0.01. (E) Behavioral entropy, measured as MoSeq syllable transition matrix entropy, was increased in NMDAR-immunized mice on clozapine compared to control diet. (F) Novel object recognition index did not significantly differ between NMDAR-immunized mice fed clozapine or control diet. (G) Prepulse inhibition did not significantly differ between NMDAR-immunized mice fed clozapine or control diet. (H) Anti-NMDAR antibody concentration was reduced in NMDAR-immunized mice on clozapine compared to control diet. Dots represent individual animals. Shaded area: interquartile range. Horizontal bar: median. Group comparison by Wilcoxon’s rank sum test. (I) Inverse relation between serum clozapine levels and anti-NMDAR antibody concentration at trend level. Dots: individual animals. Dashed line: least-square regression. Statistical test: Pearson correlation. (J) - (K) Microglial expression of the IgG Fc gamma receptor FcγRI CD64 was increased in NMDAR-immunized mice on clozapine diet compared to control diet, while there was no significant difference in microglial expression of the activation marker MHC-II. All dot plots: unless otherwise specified, dots represent individual animals; horizontal line indicates group mean; shaded area indicates probability density (C, E, F, G) or ±1SD (H, J, K). Time-series panel (B): lines indicate group means; shaded areas indicate SEM. Group comparisons by Welch’s t-test throughout unless otherwise noted. n.s. p>0.1 ∗p < 0.05, ∗∗p < 0.01, ∗∗∗p < 0.001.

Notably, clozapine treatment almost completely prevented psychosis-like behavior after anti-NMDAR immunization: in the home cage environment, clozapine-treated NMDAR-immunized mice did not exhibit the hyperlocomotion observed in non-treated NMDAR-immunized mice (**Fig. 6B-C**). Consistent with this, behavioral syllable analyses with MoSeq indicated that clozapine broadened of the constrained behavioral repertoire induced by anti-NMDAR immunization: clozapine-treated compared to non-treated NMDAR-immunized mice showed an increase in exploratory syllables and decrease in most stereotypical syllables (**Fig 6D**), and clozapine increased the entropy of the behavioral sequences (**Fig. 6E**). Clozapine showed less pronounced effects on cognitive and sensorimotor function: it numerically improved novel object recognition index, although this effect did not reach significance (**Fig. 6F**) and had no effect on prepulse inhibition (**Fig. 6G**).

This notwithstanding, the data taken together establish that clozapine protects against many of the psychosis-like behaviors induced by anti-NMDAR autoimmunity.

### Clozapine has immunomodulatory effects

Having established that clozapine prevents psychosis-like behaviors, we next interrogated the underlying mechanisms. Based on clinical observations of a reduction of immunoglobulin levels during clozapine treatment,^27^ we speculated that clozapine may attenuate the autoantibody response. Strikingly, clozapine-treated NMDAR-immunized mice showed reduced serum anti-NMDAR autoantibody levels compared to untreated NMDAR-immunized mice (**Fig. 6H**), and serum anti-NMDAR autoantibody levels were inversely related to serum levels of clozapine at trend level (**Fig. 6I**), indicating a dose-dependent relation between clozapine exposure and autoantibody response. We next asked whether this translated to down-stream effects on microglia. Indeed, clozapine-treated NMDAR-immunized mice showed increased microglial expression of surface CD64 compared to untreated NMDAR-immunized mice (**Fig. 6J**), consistent with decreased antibody-mediated NMDAR phagocytosis. Microglial MHC-II expression, in contrast, was unchanged in clozapine-treated compared to untreated NMDAR-immunized mice (**Fig. 6K**) suggesting that the rescue of surface CD64 was due to antibody-dependent phagocytosis rather than due to a global increase in microglial reactivity. Hence, clozapine reduced both anti-NMDAR autoantibody levels and microglial downstream effects.

The observation of clozapine’s effects on anti-NMDAR autoantibody levels and effects was unexpected and prompted us to ask whether this reflects a broader immunomodulatory property of clozapine. To test this idea, we assessed the effects of clozapine in *kika* mice, a genetic mouse model of systemic lupus erythematosus (SLE).^29^ *Kika* mice carry a human disease-associated gain-of-function mutation in the toll-like receptor 7 (TLR7-Y264H) that increases affinity for endogenous TLR7 ligands, leading to a breach of B cell tolerance with spontaneous production of anti-nucleic acid autoantibodies, thrombocytopenia and splenomegaly similar to what is observed in human SLE patients. From six weeks of age, *kika* mice were titrated onto a diet containing 750ppm clozapine and serum and spleens were collected at 16 ± 3 (mean ± SEM) weeks of age (**Fig. 7A**). As expected, untreated *kika* mice showed substantial splenomegaly compared to wild-type mice, apparent on visual inspection (**Fig. 7B**) and reflected by an approximately 5-fold increase in spleen weight (**Fig. 7C**). Notably, splenomegaly was attenuated in clozapine-treated *kika* mice (**Fig 7B-C**), indicating that clozapine ameliorated lupus-like phenotypes. A similar pattern was observed for serum autoantibodies: both anti-DNA and anti-RNA autoantibodies were elevated in untreated *kika* mice relative to wild-type mice, and reduced in clozapine-treated versus untreated *kika* mice (**Fig. 7D-E**). Hence, clozapine’s immunomodulatory effects on anti-NMDAR autoimmunity extended to SLE-like autoimmunity.

**Figure 7:**
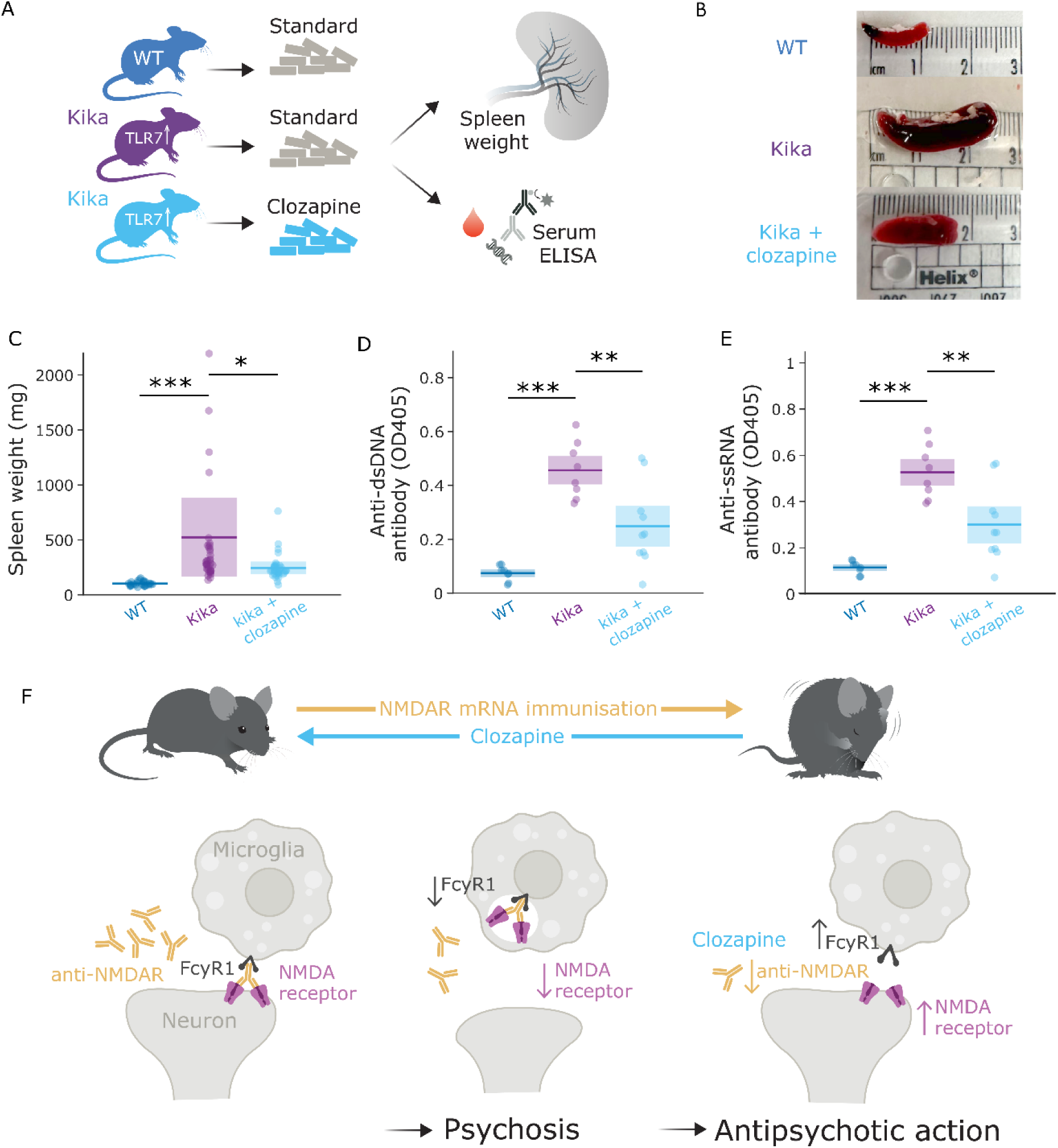
Clozapine mitigates systemic autoimmunity in a mouse model of lupus. (A) Experimental design. Kika mice, which carry a gain-of-function TLR7 mutation (TLR7-Y264H) associated with human lupus, weaned on to a diet containing 750ppm clozapine starting at 6 weeks of age. Serum antibody levels and spleen weight were assessed … (B) Representative spleen samples from wild-type and kika mice on standard diet and kika mice on clozapine diet collected at endpoint (15 weeks of age). (C) Splenomegaly in kika mice was reduced by clozapine treatment. (D) - (E) Anti-nucleic acid antibodies in serum of kika mice were reduced by clozapine treatment, measured as optical density from ELISAs for anti-dsDNA antibodies (D) and anti-ssRNA antibodies (E) All dot plots: dots represent individual animals; horizontal line indicates group mean; shaded area indicates ±1SD. Group comparisons by pairwise Welch’s t-tests. ∗p < 0.05, ∗∗p < 0.01, ∗∗∗p < 0.001. (F) Graphical Abstract - schematic of proposed mechanism. Upper panel: NMDAR mRNA immunization induces psychosis-like state, which is suppressed by clozapine. Lower panel: anti-NMDAR antibodies bind to neuronal NMDAR triggering microglial phagocytosis, leading to NMDAR loss and psychosis. Clozapine reduces anti-NMDAR antibody levels, thereby preventing NMDAR phagocytosis, rescuing NMDAR levels and producing anti-psychotic effects.

In summary, these results demonstrate that clozapine suppresses autoantibody responses across mechanistically distinct models of autoimmunity.

## DISCUSSION

Here, we report a novel mouse model of acute psychosis that uses mRNA-LNP immunization against the NMDAR to recapitulate key behavioral features of psychosis. Mechanistically, we show that psychosis-like behavior is caused by anti-NMDAR autoantibodies, accompanied by antibody-dependent NMDAR phagocytosis in microglia and loss of NMDAR (**Fig. 7F**). Using this model, we demonstrate that the most effective antipsychotic drug, clozapine, prevents psychosis-like behavior at least in part through suppression of the autoantibody response - an unexpected finding that we replicate in a mechanistically distinct model of systemic autoimmunity - suggesting that immunomodulation may be a previously unrecognized contributor to clozapine’s unique antipsychotic efficacy.

Our work extends previous mechanistic work on anti-NMDAR autoimmunity by demonstrating a causal contribution of antibody-dependent NMDAR phagocytosis to psychosis-like behavior. Both passive transfer models using patient-derived autoantibodies and active immunization models have established a link between anti-NMDAR autoantibodies and behavioral phenotypes,^30^ yet the underlying effector mechanisms have remained unclear. Notably, a recent active immunization study reports microglial activation and NMDAR-IgG complexes within microglial endosomes,^24^ providing correlative evidence that microglia participates in removal of antibody-NMDAR complexes. Here, we confirm these findings through convergent evidence from immunofluorescence microscopy, immunoblotting, and immuno-electron microscopy, collectively demonstrating NMDAR loss and accumulation of NMDAR within microglial endosomes. We extend these observations through flow cytometry evidence of microglial FcyRI internalization, implicating antibody-dependent phagocytosis as the underlying effector mechanism. Crucially, we establish the causal contribution of this mechanism through the observation that FcεRIγ-knockout mice that lack the full molecular machinery for antibody-mediated phagocytosis show attenuated behavioral phenotypes, despite comparable levels of autoantibodies. Hence, our findings support a model in which anti-NMDAR autoantibodies engage microglial FcγR, leading to engulfment of NMDAR-IgG complexes, with this antibody-dependent phagocytosis driving overall NMDAR loss and psychosis-like behavior.

Our unexpected finding that clozapine suppresses autoantibody responses, which in our model of acute psychosis translated to reduced antibody-dependent phagocytosis and attenuated psychosis-like behavior, raises the possibility that immunomodulation may contribute to clozapine’s particularly potent antipsychotic efficacy. Indeed, reductions of total IgG levels have been observed in schizophrenia patients treated with clozapine,^28^ suggesting that the immunomodulatory effects we observe in our mouse models are conserved in humans. However, whether these immunomodulatory effects contribute to antipsychotic action in schizophrenia patients remains to be established. While our mouse model of acute psychosis shares important features associated with schizophrenia such as NMDAR hypofunction and psychosis-like behaviors, it recapitulates anti-NMDAR encephalitis, a disease that is distinct from schizophrenia and has a clear autoimmune pathogenesis. Nevertheless, genetic, epidemiological and biomarker studies implicate dysregulated immune responses in schizophrenia.^31^ Moreover, subgroups of patients with schizophrenia or related conditions have brain-binding antibodies in their CSF^32^ or lymphocyte infiltrations in the brain,^33,34^ consistent with a brain autoimmune process underlying psychotic symptoms in a subset of schizophrenia patients. This raises the intriguing possibility that clozapine’s immunomodulatory properties may be the mechanism of antipsychotic action in this immune-dysregulated subset of schizophrenia patients, potentially explaining its well-documented efficacy in patients who have failed other treatments.^35^

Overall, our work establishes a mouse model of acute psychosis that implicates antibody-dependent NMDAR phagocytosis as a causal driver of psychosis-like behavior in anti-NMDAR autoimmunity and uncovers a previously unrecognized immunomodulatory mechanism of antipsychotic action, opening new avenues for targeted treatments for schizophrenia.

## RESOURCE AVAILABILITY

### Lead contact

Further information and requests for resources and reagents should be directed to and will be fulfilled by the lead contact, Katharina Schmack

### Materials availability

All unique/stable reagents generated in this study are available from the lead contact with a completed materials transfer agreement.

### Data and code availability

All raw data supporting the findings of this study is available on Figshare at https://doi.org/10.25418/crick.31333318. All analysis code required to reproduce the figures is available on Github at https://github.com/FrancisCrickInstitute/HeFeldman2026. Both repositories will be made publicly accessible upon publication.

## ACKNOWLEDGEMENTS

This work was supported by the Wellcome Trust (reference 226779/Z/22/Z) and by the Francis Crick Institute which receives its core funding from Cancer Research UK (CC2222), the UK Medical Research Council (CC2222), and the Wellcome Trust (CC2222). Harriet Feldman is a recipient of a NARSAD Young Investigator Award from the Brain & Behavior Research Foundation (grant ID 31634). Antonia Landwehr received an award provided by the BranchOut Neurological Foundation. Marie-Ève Tremblay is a Canada Research Chair (Tier I) in Neurobiology of Healthy Cognitive Aging (CRC-2024-00155). The Tremblay Lab’s Zeiss Crossbeam 350 scanning electron microscope was acquired with funding from a Canada Foundation for Innovation John R. Evans Leaders Fund grant (39965 Laboratory of ultrastructural insights into the neurobiology of aging and cognition).

We thank Sara Salgueiro Torres, Stefania Marcotti, and the Advanced Light Microscopy STP at the Francis Crick Institute for providing equipment and advice for the immunofluorescence experiments. We thank Gavin Kelly, Kanad Mandke, and Lina Gerontogianni from the Bioinformatics and Statistical STP for statistical advice on immunological and behavioral data analysis. We are grateful to Caroline Zverev, Bobbi Clayton, Ryan Hoskins, Joy Taylor, Trinity Camacho, Harry Glen, Matthew Knowles, Arturo Fernandez and other members of the Biological Research Facility STP at the Francis Crick Institute for animal provision, animal care, assistance with behavioral experiments and veterinary advice on animal welfare. We thank the In-Vivo Imaging team of the Biological Research Facility at the Francis Crick Institute for provision of equipment for validation experiments. We thank Mary Green, Ania Mikolajczak, and the Experimental Histopathology STP at the Francis Crick Institute for assistance with FFPE staining for immunofluorescence experiments. We thank Adam Hurst from the Mechanical Engineering Workshop at the Francis Crick Institute for building the behavioral setups for the MoSeq experiments. We thank the Flow Cytometry STP at the Francis Crick Institute for providing equipment and technical assistance for the flow cytometry experiments. We thank James MacRae and the Metabolomics STP at the Francis Crick Institute for providing support for the clozapine serum level analysis. We thank the Scientific Computing STP at the Francis Crick Institute for providing server space and computing resources for data analysis. We thank Chloe Roustan from the Structure Biology STP at the Francis Crick Institute for assistance with producing the NMDAR antigen and antibody. We thank Molly Strom of the Vector Core at the Francis Crick Institute for assistance with plasmid vector design. We thank Colin Murray at the School of Medical Sciences, Faculty of Health, University of Victoria, for assisting with the immunoperoxidase staining. We thank Caetano Reis e Sousa, Oliver Schulz, Bradley Jamieson, Karl Brune, Lukasz Wieteska, Lingling Zhang and Mathieu Bourdenx for valuable intellectual input and experimental suggestions. Julia Kuhl created the schematic illustrations used in the figures.

## AUTHOR CONTRIBUTIONS

Conceptualization: LH, HF, TN, JM, and KS; methodology: LH, HF, TN, MB, VP, AB, SK, and EMT; software: LH, HF, and MB; validation: LH, HF, TN, AB, and MP; formal analysis: LH, HF, TN, MB, EMT, and AL; investigation: LH, TN, MB, VP, AB, OK, JZ, AL, and MK; resources: SK, EMT, CV, AH, and KS; data curation: LH, HF, and MB; writing – original draft: LH, HF, and KS; writing – review & editing: all authors; visualization: LH, HF, VP, and AL; supervision: SK, JM, TP, and KS; project administration: MB and KS; funding acquisition: HF, AH, and KS.

## DECLARATION OF INTEREST

Le He and Katharina Schmack are co-inventors on a patent application describing NMDAR constructs (UK Patent Application No 2606937.7). Thomas A. Pollak has received consulting fees from Arialys Therapeutics.

## STAR★METHODS

### KEY RESOURCES TABLE

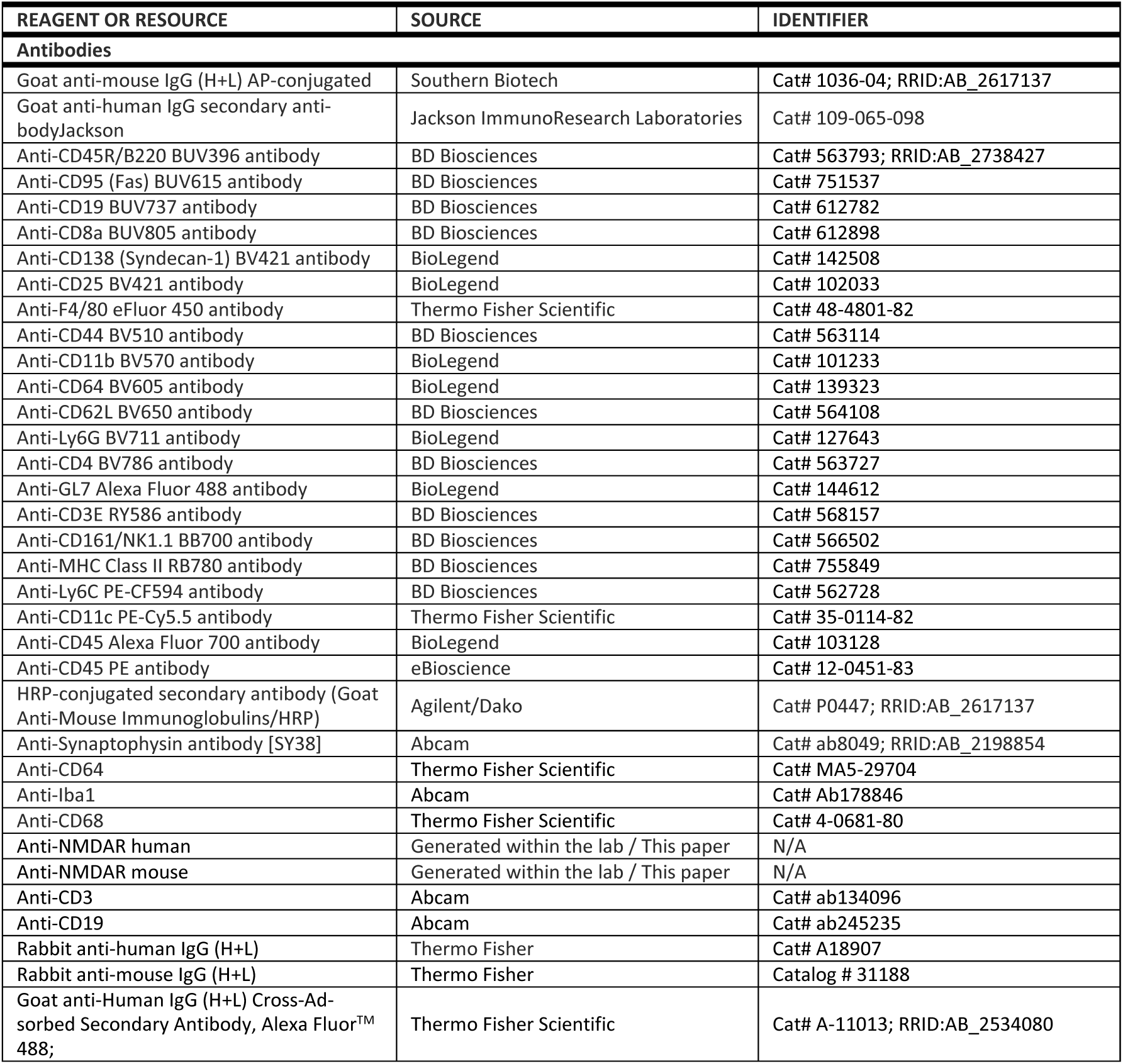
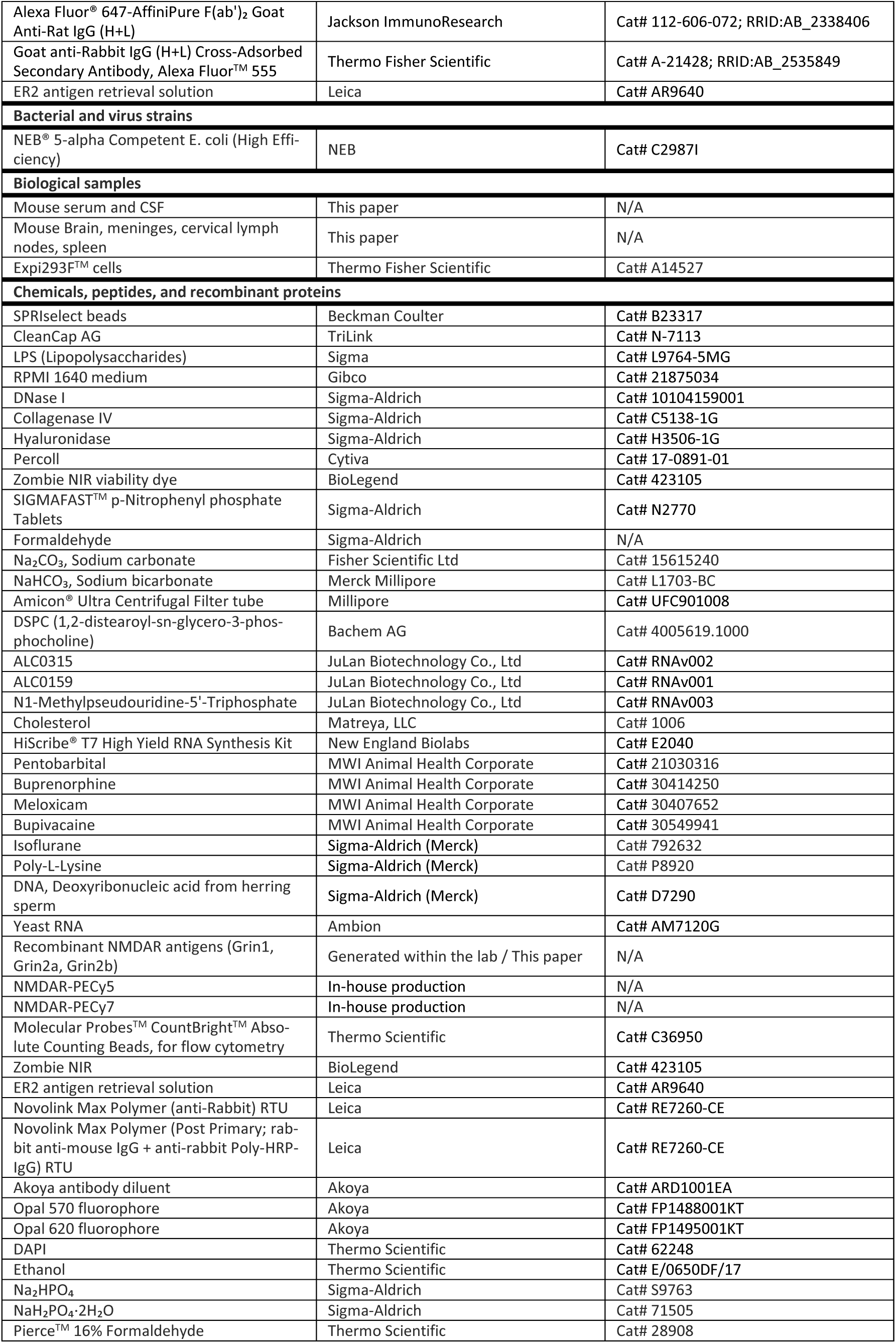
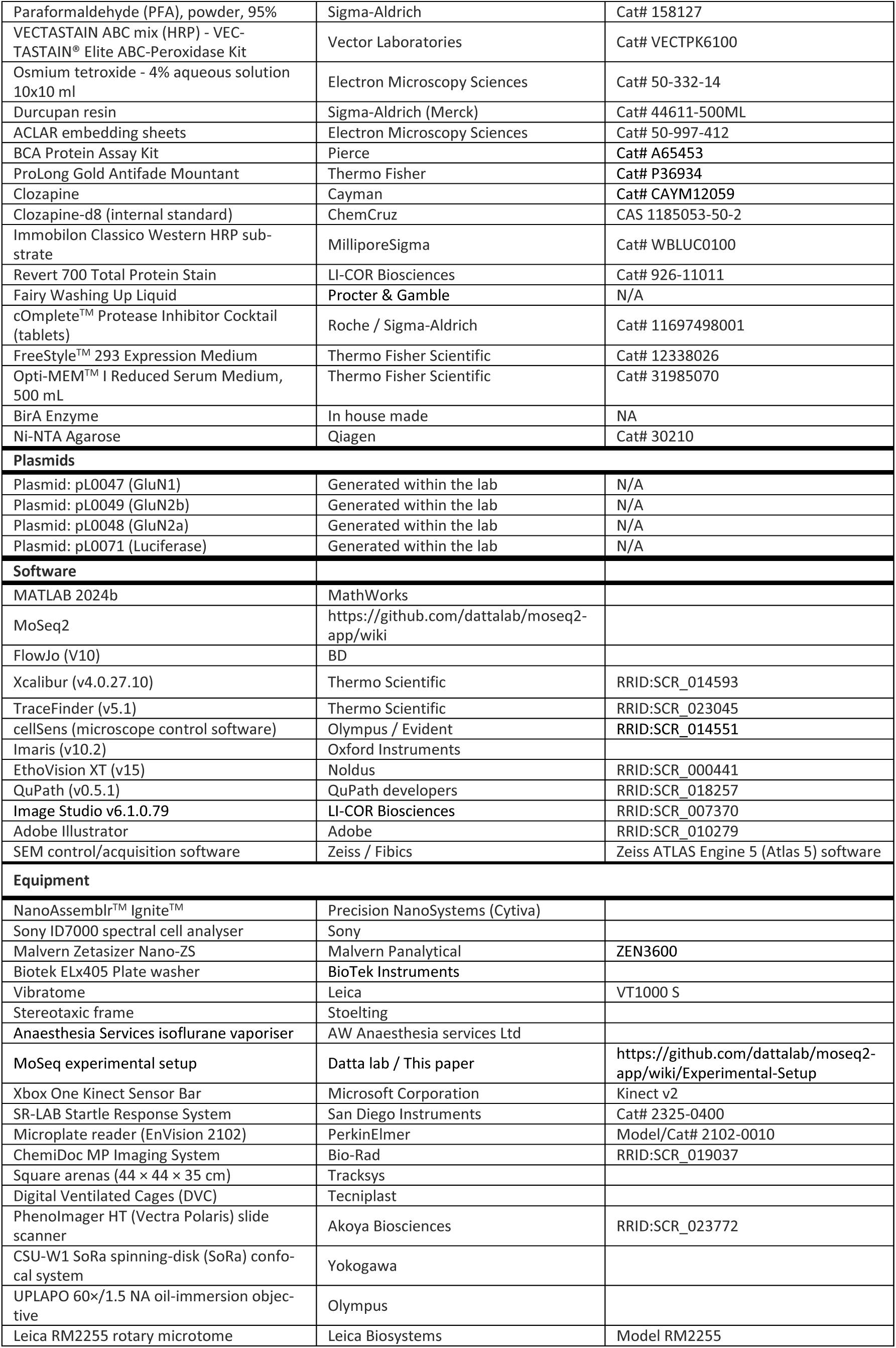
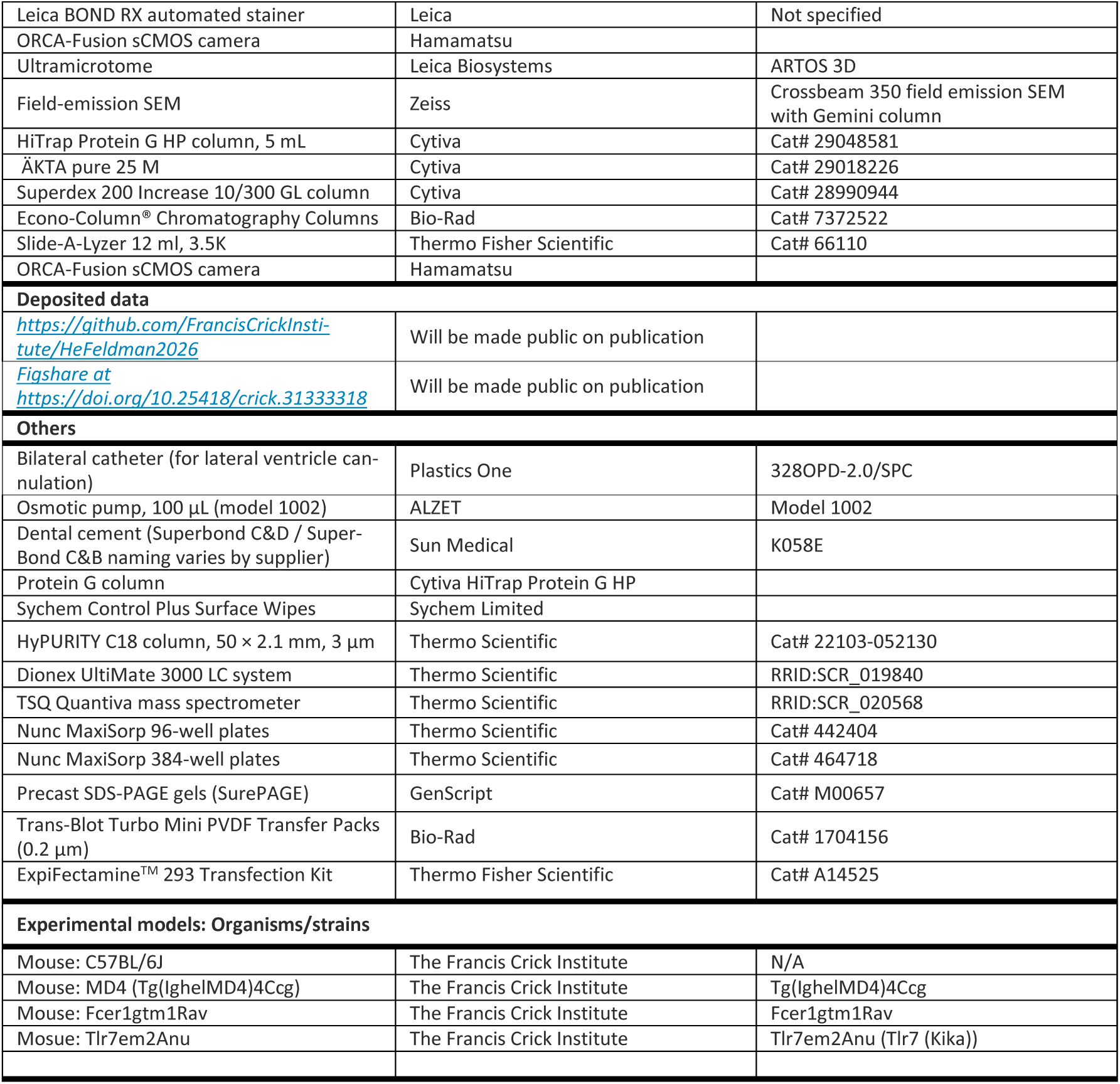

### EXPERIMENTAL MODEL

#### Animals

Adult C57BL/6J mice, MD4 transgenic mice (Tg(IghelMD4)4Ccg), Fcer1g knockout mice (Fcer1gtm1Rav) and TLR7 kika mice (Tlr7em2Anu) were used in this study. All mice were older than 6 weeks at the start of the experiments. Both male and female mice were included. Animals were housed in groups of maximum 5 per cage under standard conditions (12-hour light/dark cycle, temperature 22 °C, humidity 50%) with food and water available ad libitum. For experiments involving Digital Ventilated Cages (DVC**®**, Tecniplast), mice were single-housed.

For behavioral experiments, mice were transported to the behavioral rooms and habituated there for 0.5 hours before each testing session to minimize stress related to handling and environmental change. All animal-related procedures were performed in accordance with national and institutional guidelines for animal care and were approved by the Francis Crick Institute - Biological Resources Facility Strategic Oversight Committee (incorporating the Animal Welfare and Ethical Review Body) and by the Home Office, UK (Project licence PP2867252).

#### mRNA-LNP formulation and dosing for anti-NMDAR immunization

To induce an autoimmune response against the NMDAR, we used mRNA encapsulated in lipid nanoparticles (mRNA-LNP). This delivery platform enables efficient in-vivo expression of the antigen encoded in the mRNA, thereby eliciting robust immune responses against that antigen.

mRNA-LNP formulations were adapted from the Pfizer-BioNTech BNT162b2 mRNA Covid-19 vaccine,^36^ using an ethanol-dissolved lipid mixture composed of ALC-0315, DSPC, cholesterol, and ALC-0159 at a molar ratio of 46.3:9.4:42.7:1.6. ^36^ Plasmids encoding the NMDAR subunits GluN1 (pL0047), GluN2A (pL0048), GluN2B (pL0049), or luciferase were generated in-house, linerized with SpeI and SnaBI, and purified using SPRIselect beads (B23317, Beckman Coulter) prior to in vitro transcription. mRNA was synthesized using the HiScribe T7 High Yield RNA Synthesis Kit (E2040, New England Biolabs) with complete substitution of uridine-5′-triphosphate by N1-Methylpseudouridine-5’-Triphosphate and incorporation of CleanCap AG (N-7113, TriLink) according to the manufacturer’s instructions in 40 µL reactions. For NMDAR constructs, GluN1, GluN2A, and GluN2B mRNAs were combined at a 1:1:1 weight ratio; NMDAR and luciferase mRNAs were then mixed with the lipid solution at an N/P ratio of 6 using a NanoAssemblr Ignite microfluidic mixer (Precision NanoSystems) to generate mRNA-LNPs.

The crude mRNA-LNP suspension was purified using Amicon Ultra centrifugal filter units (UFC901008, Millipore) to remove ethanol, and the retentate was resuspended in 10 mM Tris-HCl containing 20 mg/mL sucrose to a final concentration of 0.1 mg mRNA/mL. Particle size was measured using a Malvern Zetasizer Nano-ZS (Malvern Panalytical), and only preparations with a mean hydrodynamic diameter <200 nm were used for in vivo experiments. Aliquots were stored at −80 °C and thawed immediately before injection, avoiding repeated freeze-thaw cycles.

On each immunization, mice received 100 µL of mRNA-LNP suspension containing 10 µg mRNA, co-administered with 2 µg lipopolysaccharide (LPS; L9764-5MG, Sigma; stock solution diluted in sterile H2O) by subcutaneous injection. Immunization sites (e.g., head/back flank or scruff) were alternated between doses to reduce local skin irritation. Despite site rotation, visible injection-site lesions occurred in around 10% of mice, and such animals were monitored closely and managed according to predefined humane endpoints.

Immunizations were administered at two-week intervals. The number of immunizations varied by experimental cohort according to the scientific objective: mice designated for behavioral phenotyping received three immunizations, allowing behavioral assessments to be performed between and after immunization rounds to capture the emergence and progression of psychosis-like behavior over time. Mice designated for immune phenotyping received two immunizations and were culled one week after the second immunization, a timepoint chosen to capture the immune phenotype at the onset of behavioral changes observed in the behavioral cohort. The cohort in Figure 5 followed the behavioral protocol but the third immunization was delayed by one week because behavioral differences between wild-type and Fcer1g knockout mice were already apparent following the second immunization; extending the interval allowed us to assess whether the between-group divergence continued to expand prior to the final immunization.

### BEHAVIORAL PHENOTYPING

#### Home cage activity monitoring

To assess home cage activity, mice were transferred to single housing in Digital Ventilated Cages (DVC) starting a few days prior to the first immunization. These cages continuously capture animal locomotion via a grid of capacitive sensors embedded in the cage floor, which detect the position and movement of the mice.

Data were collected continuously throughout the entire experimental period. Home cage monitoring was performed for all experimental groups from at least 4 days before the first immunization to week 7 after the first immunization. Home cage data for the passive transfer experiment (Fig.3) was dominated by individual differences in differential impact of the minipumps on movement restriction and is therefore not presented.

#### Motion sequencing (MoSeq)

To assess the structure of spontaneous behavior, we used Motion Sequencing (MoSeq), a machine learning-based approach that uses depth imaging to decompose mouse behavior into a repertoire of short movement segments called **“**syllables”. During data acquisition, mice are recorded from above using a depth camera as they freely explore an open arena. The depth data captures a 3D representation of the animal’s posture at each frame. MoSeq then applies an autoregressive hidden Markov model to segment continuous movement into discrete, recurring postural motifs, typically lasting 350 - 500 ms, that together constitute the animal’s behavioral repertoire.

Experimental sessions lasted 30 minutes and were performed in black high-walled circular arenas (43 cm diameter), as described previously (https://github.com/dattalab/moseq2-app/wiki#hardware-setup-for-data-acquisition).^37^ Data for MoSeq analysis were captured using a Kinect infra-red depth camera. The day before the first data acquisition session, mice were habituated for 10min in the arena. During the mRNA immunization experiments, data quality of video recordings from one of the four arenas was insufficient to allow for behavioral syllable analyses, which meant that these sessions were excluded from MoSeq analyses. However, mouse position could still be accurately inferred, therefore these sessions were included into open field analysis.

MoSeq was performed for the mRNA immunization (Fig. 1), passive transfer (Fig. 3) and clozapine (Fig. 6) experiments. In the mRNA immunization and clozapine experiments, MoSeq data from open field tests were collected at weeks –1, 3, 5 and 7 with respect to the first immunization. In addition, novel object recognition data was collected in the MoSeq arenas and the video data analyzed with MoSeq. In the passive immunization experiments, data were collected at days 5, 9 and 14 after implantation.

#### Novel Object Recognition

To assess cognitive function, we used the Novel Object Recognition assay, which is widely used in rodents to assess learning and memory. The assay measures how much time a mouse spends exploring a novel object versus a familiar one, exploiting the innate preference of rodents for novelty.

The assay consisted of three sessions: a habituation session, a familiarization session, and a test session. During the habituation session, conducted one week before the other sessions, mice were placed in an empty arena for 30 minutes. During the familiarization session, mice were first placed in the open-field arena containing two identical objects for 10 minutes. During the test sessions, conducted three hours after the familiarization session, mice were then returned to the same arena for another 10 minutes session, during which one of the familiar objects was replaced with a novel object. Between each mouse, the arena was cleaned with Sychem surface wipes.

Novel object recognition was tested for the mRNA immunization (Fig. 1), passive transfer (Fig. 3) and clozapine (Fig. 6) experiments. For the mRNA immunization experiment, data was collected from one familiarization and one testing session in week 7. For the passive transfer experiment, a familiarization and test session was conducted on days 4, 9 and 13.

The assay for the passive transfer experiment (Figure 2) was conducted in four square arenas (44 × 44 × 35 cm; Tracksys) positioned on an illuminated platform, whereas the assay for mRNA immunization experiment (Figures 1 and 6) was performed in the four circular MoSeq arenas (diameter 42 cm, height 36 cm). All trials were video-recorded, and EthoVision XT (v15, Noldus) was used to track mouse position within the arena and predefined object zones. Raw data were exported and analyzed using custom MATLAB code (MathWorks, R2024b).

#### Prepulse Inhibition

To assess sensorimotor gating deficits characteristic of schizophrenia, we used the prepulse inhibition assay, a measure the reduction in the magnitude of the acoustic startle response when a weak, non-startling sound - the prepulse - precedes an intense, potentially startling, sound. All testing was conducted using SR-LAB-Startle Response System (SanDiego Instrument) under continuous background noise (70 dB). Prepulse inhibition was measured for the mRNA immunization (Fig. 1) and clozapine (Fig. 6) experiments. Prepulse inhibition was not performed for the passive transfer experiment because of the risk of displacement of the osmotic minipumps in the startle chamber.

One day prior to testing, mice were habituated to the apparatus for 10 minutes. During the experimental session, a range of stimulus intensities (25-50dB above background) was used to characterize the full psychometric startle response curve. Prepulse trials consisted of 20ms of white noise (16 dB above background), followed 50ms later by the startle stimulus. Trials were presented both with and without the prepulse across all stimulus intensities, allowing the prepulse inhibition to be calculated as the percentage reduction in startle amplitude on prepulse trials relative to no-prepulse trials. Between each mouse, the apparatus was cleaned with Sychem surface wipes.

Prepulse inhibition data was collected at weeks –1, 3 and 5 of the mRNA immunization experiment with respect to the first immunization.

### ANTIBODY PROFILING AND PASSIVE TRANSFER

#### Serum collection

Serum was collected for quantification of autoantibody concentration, IgG purification for passive transfer experiments and for clozapine concentration measures. Depending on the timepoint, serum was collected either by lateral saphenous vein bleed during the experimental period or by cardiac puncture under deep anesthesia after a lethal dose of pentobarbital at the experimental endpoint.

#### Cerebrospinal fluid collection

To measure CNS autoantibodies, cerebrospinal fluid (CSF) was collected from anaesthetized mice following a modified version of the protocol described previously.^38^

Animals were anaesthetized under isoflurane (5% induction, 2–2.5% maintenance) in oxygen. Following loss of pedal reflex, the mouse was transferred to a stereotaxic frame (Stoelting) equipped with a temperature-controlled heating pad to maintain normothermia. The dorsal neck region and occipital area were shaved, cleaned with 0.05% chlorhexidine or sterile saline, and exposed surgically to visualize the cisterna magna. Using fine scissors, the overlying connective tissue and muscles were gently separated along the midline to avoid bleeding and reveal the translucent cisternal membrane.

A pulled glass capillary attached via silicone tubing to a 20 mL syringe was positioned horizontally with a micromanipulator. Under magnification, the capillary tip was advanced toward the cisterna magna, avoiding visible blood vessels, until slight resistance indicated contact with the membrane. The membrane was carefully punctured, allowing clear CSF to enter the capillary by capillary action. Between 8-15 µL of CSF was collected over approximately 10 min. Samples were expelled into pre-labelled low-protein-binding microcentrifuge tubes by gentle air pressure from the syringe and immediately stored on ice before subsequent processing. At the end of the procedure, animals were euthanized.

#### Enzyme-linked immunosorbent assay (ELISA) for anti-NMDAR antibodies

As no commercial ELISA exists for the detection of anti-NMDAR IgG antibodies; we devised a novel assay. The assay uses a recombinant NMDAR chimeric protein as the capture antigen and an in-house-produced anti-NMDAR monoclonal IgG antibody to generate a standard curve for quantification. The following sections describe the production and validation of each reagent, followed by the full ELISA protocol.

##### NMDAR chimeric antigen production

To produce the ELISA capture antigen, a plasmid encoding a chimeric NMDAR extracellular domain construct was generated in-house. The construct comprised GluN1 and GluN2A or Grin2B extracellular domain, fused to a C-terminal hexahistidine (6 x His) tag for affinity purification and an AviTag sequence (GLNDIFEAQKIEWHE) for site-specific biotinylation. The AviTag serves as the sole substrate for the bacterial biotin ligase BirA, enabling covalent attachment of a single biotin molecule to a defined lysine residue within the tag; encoding the AviTag within the plasmid construct restricts biotinylation to this site and thereby preserves the three-dimensional conformation of the antigen.

To produce recombinant NMDAR chimeric antigen for use as the ELISA capture antigen, Expi293F^TM^ cells (Thermo Fisher Scientific) were transiently transfected with the antigen-encoding plasmid in a 1 L format. Cells were cultured in Expi293^TM^ Expression Medium at 37°C, 8% CO₂, 125–130 rpm to a density of 3 × 10⁶ cells/mL (>95% viability). Plasmid DNA (1 mg/L total) and ExpiFectamine^TM^ 293 reagent (3:1 vol:vol reagent:DNA) were each diluted separately in OptiPRO^TM^ SFM, combined, incubated for 20 min at room temperature, and added directly to cells. Approximately 20 h post-transfection, ExpiFectamine^TM^ Enhancer 1 (1 mL per 200 mL culture) and Enhancer 2 (10 mL per 200 mL culture) were added without media change. Cultures were shifted to 32°C following enhancer addition to improve folding of the secreted chimeric antigen. Supernatants were harvested 4 days post-transfection by centrifugation (300-500 × g, 5 min), clarified (1,500 × g, 20 min), and sterile-filtered (0.22 μm).

Filtered supernatant was incubated with NiNTA beads equilibrated in PBS. The bead-supernatant mixture was applied to an empty Bio-Rad gravity column, washed with 10 mM imidazole in PBS, and His-tagged antigen was eluted with 300 mM imidazole in PBS. Eluate was dialysed twice against PBS (10 kDa MWCO Slide-A-Lyzer, 2.5 L per dialysis, 4°C) to remove imidazole.

Site-specific biotinylation was performed to enable streptavidin-based capture of the antigen in downstream applications. Chemical biotinylation reagents (e.g., NHS-biotin) non-selectively modify surface-exposed lysine residues, which may include residues within or adjacent to antibody-binding epitopes and can thereby alter antigen conformation and reduce assay sensitivity. To avoid this, the antigen construct was designed to carry an AviTag sequence (see Plasmid design above), which restricts biotinylation by BirA enzyme to a single, defined site remote from the antigen surface.

Reactions were assembled on ice with the following final concentrations: 5 mM MgCl2, 5 mM ATP (pH 8.0), 0.3 mM D-biotin (pH 8.0), and a target protein:BirA molar ratio of 50:1, using in-house-produced BirA enzyme. Reactions were incubated overnight at 4°C. Biotinylated antigen was subsequently purified by size-exclusion chromatography on a Superdex 200 Increase column (Cytiva) equilibrated in PBS to remove free biotin and reaction components. Biotinylation efficiency was confirmed by streptavidin bead pull-down followed by western blot using streptavidin-HRP, which produces signal only in the presence of successful biotin conjugation.

Purified antigen was aliquoted, quantified by absorbance at A280, sterile-filtered (0.22 μm), and stored at −80°C.

##### Anti-NMDAR IgG antibody production

A recombinant anti-NMDAR monoclonal IgG antibody was produced in-house for use as the quantification standard in the ELISA and also imaging experiments (see below). Two plasmids were constructed, each encoding the variable region of the validated anti-NMDAR antibody N308/48^39^: one with a mouse IgG2a constant region for use as the ELISA quantification standard, and one with a human IgG1 constant region for use in imaging experiments. Both were used for expression.

Expression was carried out in Expi293F™ cells by transient transfection with the antibody-encoding plasmid, following the same conditions described above for antigen production, with the exception that cultures were maintained at 37°C throughout (rather than being shifted to 32°C). Supernatants were harvested and clarified as described above.

Filtered supernatant was loaded onto a 5 mL HiTrap Protein G column (Cytiva) equilibrated in 0.1 M Tris (pH 8) at 1–5 mL/min using an ÄKTA system. The column was washed with 10 column volumes of PBS to baseline A280, and IgG was eluted with 0.1 M glycine (pH 3.0) directly into tubes pre-containing an equal volume of 1 M Tris-HCl (pH 8.0) for immediate neutralisation. Peak fractions (monitored at A280) were pooled and further purified by size-exclusion chromatography on a Superdex 200 Increase column (Cytiva) equilibrated in PBS.

Purified antibody was aliquoted, quantified by absorbance at A280, sterile-filtered (0.22 μm), and stored at −80°C.

##### ELISA for anti-NMDAR autoantibodies measurement

We used the recombinant NMDAR antigen and anti-NMDAR antibody to design and validate an ELISA to detect IgG antibodies specific for the NMDAR.

MaxiSorp plates (96- or 384-well) were coated overnight at 4 °C with recombinant NMDAR antigens (5 µg/mL, 40uL or 15uL) in carbonate-bicarbonate buffer (1.95 g Na₂CO₃ and 2.93 g NaHCO₃ in 1 L ddH₂O, pH 9.6). Plates were washed four times with PBS containing 0.05% Tween-20 (PBST) using a Biotek plate washer. To minimize nonspecific binding, wells were blocked with 2% goat serum in PBS (blocking buffer) for 1 hour at room temperature, followed by an additional four PBS-T washes. Serial dilutions of in-house-produced anti-NMDAR mouse IgG antibody, as well as serum diluted 1:20 in blocking buffer, and cerebrospinal fluid (CSF) samples, diluted 1:3, were incubated in the antigen-coated wells for 1 hour at room temperature. After removal of unbound materials by four PBST washes, goat anti-mouse IgG (H+L) alkaline phosphatase-conjugated antibody (Southern Biotech, Cat. No. 1036-04; 1:2,500 dilution in PBS) was added and incubated for 1 hour at room temperature. The plates were then washed four times with PBS-T. Colorimetric signal was developed using SIGMAFAST^TM^ p-Nitrophenyl phosphate Tablets (N2770, 50 tests, Sigma-Aldrich), prepared according to the manufacturer’s protocol, and optical density was measured at 405nm using the EnVision 2102 Multilabel Plate Reader, model 2102-0010 (PerkinElmer).

#### Passive transfer of IgG into the cerebral ventricle via osmotic pumps

To directly assess whether autoantibodies were sufficient to induce behavioral phenotypes, purified IgG from immunized donors was infused into the cerebral ventricles of naïve recipient mice via implanted osmotic pumps, enabling continuous intracerebroventricular delivery over weeks.

IgG antibodies were collected from the serum of mice immunized with either NMDAR or luciferase mRNA-LNP. Animals were monitored using Digital Ventilated Cages (DVC; Tecniplast and Olden Labs) and daily health checks and were exsanguinated upon onset of an abnormal phenotype (NMDAR group) or at a matched time point (luciferase cohort). IgG was purified using a Protein G column and concentrated at 15 mg/mL prior to infusion into osmotic pumps. Only samples from mice exhibiting behavioral phenotypes and NMDAR-specific IgG, as confirmed by in-house ELISA, were included.

Bilateral cannulation of the lateral ventricles was performed under isoflurane anesthesia using stereotaxic surgery. Mice received pre-operative buprenorphine (0.1 mg/kg) and meloxicam (5 mg/kg) subcutaneously, with local bupivacaine applied to the scalp and ear-bar sites. A bilateral catheter (Plastics One, 328OPD-2.0/SPC) was implanted (AP −0.1 mm, ML ±1.0 mm, DV −3.0 mm) and secured with dental cement (Superbond C&D, K058E). Each catheter arm was connected to a 100 µL osmotic pump filled with the IgG solution (model 1002, Alzet) implanted subcutaneously on the back.

Post-operatively, mice received 10 µL/g warm saline and recovered in a heated chamber until ambulatory before return to home cages with wet mash food. Histological analysis confirmed correct ventricular catheter placement in all animals.

### IMMUNOPHENOTYPING

#### Flow cytometry analysis of brain immune cells

To characterize immune cell changes in the brain following immunization against NMDAR, we performed flow cytometry on dissociated brain tissue.

Mice were euthanized with a lethal dose of pentobarbital (intraperitoneal injection). Under deep anesthesia, mice received an intravenous injection of anti-CD45 PE antibody (12-0451-83, eBioscience) and were rested for 5 min to label circulating immune cells. The brain was dissected and stored in PBS at 4 °C for less than 1 hour before processing. Brain tissue was incubated in 6 mL of enzyme solution; enzymatic digestion was stopped by adding EDTA to a final concentration of 1 mM. Brain-derived immune cells were enriched using 30% Percoll (17-0891-01, Cytiva) gradient centrifugation at 700 × g for 20 min at 4 °C.

Cells were stained overnight using an adapted version of a previously published protocol.^40^ Cell suspensions were stained with Zombie NIR viability dye (423105, BioLegend) to identify dead cells, then fixed with 0.2% formaldehyde (Sigma-Aldrich) for 30 min at 4 °C. Fixed cells were stained with the following panel overnight at 4 °C. To characterise immune cell infiltration and inflammation in the immunized mouse brain, cells were stained with the following antibodies: anti-CD45R/B220 BUV396, anti-CD95 (Fas) BUV615, anti-CD19 BUV737, anti-CD8a BUV805, anti-CD138 (Syndecan-1) BV421, anti-F4/80 eFluor 450, anti-CD44 BV510, anti-CD11b BV570, anti-CD64 BV605, anti-CD62L BV650, anti-Ly6G BV711, anti-CD4 BV786, anti-GL7 Alexa Fluor 488, anti-CD3ε RY586, anti-CD161/NK1.1 BB700, anti-MHC Class II RB780, anti-Ly6C PE-CF594, anti-CD11c PE-Cy5.5, anti-CD45 Alexa Fluor 700, and two in-house-produced NMDAR antigen conjugates (PE-Cy5 and PE-Cy7) for identification of NMDAR-specific B cells by dual-colour gating. Full antibody details are provided in the Key Resources Table. Following washing steps (700 × g with breaks off, 20 min, 4 °C), cells were resuspended and analyzed on a Sony ID7000 flow cytometer. For intracellular CD64 staining, the cell was further permeabilized with diluted Fairy buffer (PBS with 0.05% Fairy) at room temperature for 30min,^41^ then washed with PBS and stained with antibody for 2h at room temperature before flow cytometer analysis.

#### Formalin-fixed, paraffin-embedded (FFPE) brain tissue immunofluorescence

For quantifying IgG deposition and NMDAR levels in brain tissue, formalin-fixed, paraffin-embedded (FFPE) sections were stained for IgG and the GluN1 subunit of the NMDAR.

Brains were collected after intraperitoneal injection with a lethal dose pentobarbital and transcardial perfusion with PBS, followed by 4% PFA (Cat# 28908, diluted with PBS till 4%). The brains were dissected out from skull, fixed in 4% PFA for 24 h and stored in 70% ethanol before processing and embedding. Sections (3 µm) were cut with Leica RM 2255, baked at 60 °C for 1 h, and stained on a Leica BOND RX automated stainer as a six-panel multiplex immunofluorescence panel plus DAPI; here we report results for IgG and NMDAR. Antigen retrieval was performed before each cycle using ER2 buffer (Leica, AR9640; 95 °C for 20 min).

To visualize IgG deposition and NMDAR expression in brain tissue, two targets were stained sequentially in separate cycles using a tyramide signal amplification (TSA) multiplex protocol on the Leica BOND RX automated stainer.

The endogenous mouse IgG deposited in the brain was detected firstly. Because the Novolink Max HRP polymer recognises only rabbit IgG, a two-step bridge approach was required: a post-primary rabbit anti-mouse IgG bridge antibody (Leica Biosystems, RE7111, part of RE7260-CE; <10 μg/mL in 10% animal serum/TBS with 0.1% ProClin 950) was applied to convert the mouse-IgG signal into a rabbit-IgG-compatible substrate, followed by the anti-rabbit HRP polymer (Novolink Max, Leica, RE7260-CE). Signal was developed by tyramide signal amplification (TSA) to deposit an Opal fluorophore onto the tissue. Then, NMDAR was detected using the in-house-produced human anti-NMDAR antibody (as described in ‘ELISA for anti-NMDAR antibodies’; 1:200) in Akoya anti-body diluent. A rabbit anti-human IgG secondary antibody (Thermo Fisher Scientific, A18907; 1:500) was applied, followed by the anti-rabbit HRP polymer (anti-rabbit Poly-HRP-IgG, RTU, Novolink Max, Leica, RE7260-CE) and TSA fluorophore deposition.

Slides were counterstained with DAPI (Thermo Scientific, 62248; 1:2500) and mounted with ProLong Gold Anti-fade Mountant (Thermo Fisher, P36934). Fluorescence images of FFPE mouse brain sections were acquired using a Slide scanner PhenoImager HT Polaris (20x, Akoya).

#### Western blot of brain tissue

To complement FFPE immunostaining with an independent quantitative measure of IgG deposition and NMDAR expression, brain tissue protein levels were assessed by Western Blot

To obtain protein samples from brain tissue, mice immunized against NMDAR and healthy controls were perfused with PBS under lethal dose of pentobarbital and whole brains were rapidly dissected to obtain the cortex, hippocampus and the cerebellum on ice, with olfactory bulbs removed. Tissue was after storage at −80 °C, weighed, minced, and homogenized on ice in 0.32 M sucrose, 5 mM HEPES, pH 7.5, supplemented with complete protease inhibitor (Roche, 11697498001) at a 1:20 w/v ratio using a Dounce homogenizer.

Homogenates were centrifuged at 1,000 × g for 10 min at 4 °C to pellet nuclei, and the supernatant was subjected to a second identical centrifugation to minimize nuclear contamination. The resultant supernatant was further centrifuged at 12,000 × g for 20 min at 4 °C to generate a crude membrane pellet. Pellets were resuspended in PBS with protease inhibitors (approximately 200 µl per brain), aliquoted, stored at −80 °C, and quantified for protein concentration using BCA (Pierce, A65453) prior to Western blot analysis.

To detect protein targets in brain homogenate and crude membrane fractions, a Western blot protocol was utilized. Crude membrane or homogenate samples were adjusted to the desired concentration, mixed with SDS sample loading buffer (with reducing agent as required), heated at 95 °C for 5 min, briefly centrifuged, and 10 µg of protein per lane was loaded onto precast SDS-PAGE gels (SurePAGE GenScript, M00657) alongside a molecular weight ladder. Gels were run in standard running buffer at 120 V for approximately 75 min in ice until the tracking dye approached the bottom of the gel. Proteins were transferred from gels onto PVDF membranes using a Trans-Blot Turbo system with pre-assembled transfer stacks (BioRad 1704156) under standard conditions (25 V, 1 A, 30 min).

Membranes were then dried on filter paper for 10 min at 37 °C. After reactivating the membrane with 100% Methanol, the membranes were rehydrated in PBS for 5 min before staining with Revert total protein stain (LiCor Bio, 926-11011) for 5 min with gentle shaking. The membrane was then washed with 6.7% (v/v) glacial acetic acid, 30% (v/v) methanol, in water followed by a rinse in distilled water. The membrane was then imaged at 680 nm on a BioRad ChemiDoc system.

Membranes were then blocked in PBS containing 0.1 % Tween-20 and 3 % BSA for 2 h at room temperature to reduce nonspecific binding. Blocked membranes were incubated with primary antibodies diluted in blocking buffer, overnight at 4 °C with gentle agitation, followed by three 10 min washes in PBST. Membranes were subsequently incubated with appropriate HRP-conjugated secondary antibody (Dako, Cat# P0447) in blocking buffer for 2 h at room temperature, washed again three times for 10 min in PBST, and chemiluminescent signal was developed using Immobilon Classico substrate (WBLUC0100), with bands imaged on the BioRad Chemidoc system with chemiluminescence exposure times adjusted to prevent saturation.

#### Floating brain tissue immunofluorescence microscopy

To assess microglial morphology and NMDAR phagocytosis, free-floating brain sections were immunostained for the microglial marker Iba1, the lysosomal marker CD68, and the GluN1 subunit of the NMDAR.

Brains were collected as described for FFPE immunofluorescence microscopy, fixed overnight in 4% PFA, and stored in PBS before sectioning into 50-µm slices on a vibratome. Sections were blocked for 1-2 h at room temperature in blocking buffer (PBS, 0.3% Triton X-100, 3% BSA, 10% normal goat serum) on a rocker, then washed three times in PBS with 0.3% Triton X-100 (10 min each). Slices were incubated with anti-Iba1 (1:500, Abcam, Cat# Ab178846), anti-CD68 (1:400, Cat# 4-0681-80), and anti-GluN1 (1:200, generated in-house, human) diluted in blocking buffer overnight at 4 °C. The next day, sections were washed five times in PBS with 0.3% Triton X-100 (≥10 min each) and then incubated with fluorophore-conjugated secondary antibodies: anti-Human A488 IgG (Cat# A-11013; RRID:AB_2534080), anti-Rabbit IgG A555 (Cat# A-21428), anti-Rat IgG A647 (Cat# 112-606-072) diluted in blocking buffer overnight at 4 °C, protected from light. On Day 3, slices were washed five times in PBS with 0.3% Triton X-100, given a final PBS wash, and mounted onto glass slides in PBS before cover slipping with fluorescence-compatible mounting medium for imaging.

Images were acquired on an inverted Olympus CSU-W1 spinning disk confocal microscope equipped with a UP-LAPO 60x/1.5 NA oil-immersion objective and controlled using Olympus cellSens software. Samples were illuminated with 405, 488, 561 and 647 nm lasers, and fluorescence was detected using standard band-pass emission filters appropriate for each fluorophore. Z-stacks were acquired with a 0.5 µm step size, and exposure time of 200 ms for all channels. Laser power and detection settings were kept constant across samples. Images were recorded using a Hamamatsu ORCA-Fusion sCMOS camera, and identical acquisition parameters were applied across all samples.

#### Immunoperoxidase staining and Electron microscopy imaging

To visualize the subcellular localization of NMDARs within microglia at ultrastructural resolution, we used immunoperoxidase electron microscopy.

To obtain the brain, under lethal dose of pentobarbital, mice were transcardially perfused with ice-cold PBS followed by ice-cold 4% PFA (Cat# 158127, only for EM) in 0.1 M phosphate buffer (PB, 100 mM, pH 7.4; 11.74 g Na₂HPO₄ and 2.4 g NaH₂PO₄·H₂O in 1L water) for 10 min. Brains were extracted and post-fixed in 4% PFA in PB for 2 h at 4 °C, then sectioned into 50-µm slices on a vibratome. Sagittal brain sections containing the dorsal hippocampus (DH), Bregma between −1.5 and −3.0, were rinsed in phosphate-buffered saline (PBS), followed by an incubation with 0.3% H₂O₂ in PBS. Next, sections were quenched with 0.1% NaBH₄ for 30 min, then washed in PBS. Subsequently, sections were incubated for 1 hour at room temperature in a blocking solution consisting of 10% fetal bovine serum, 3% bovine serum albumin, and 0.01% Triton X-100 diluted in Tris-buffered saline (TBS). Sections were then incubated with primary anti-NMDA antibody [1:100] (home-made anti-body, species: human) in blocking buffer at 4°C overnight.

Following washes in TBS, sections were incubated with a goat anti-human secondary antibody [1:300] (Jackson ImmunoResearch Laboratories, Cat# 109-065-098) for 1 hour, followed by quenching in ABC Vectastain mix (1:100 in TBS; Vector Laboratories). Anti-NMDA immunolabelling was revealed using diaminobenzidine (DAB; 0.05%) and H₂O₂ (0.015%) in TBS. Following immunostaining, sections were post-fixed in 1% osmium tetroxide, embedded in thiocarbohydrazide (TCH), and incubated again in 2% osmium tetroxide. Sections were then dehydrated through increasing concentrations of ethanol and immersed in propylene oxide, before overnight incubation in Durcupan resin (Electron Microscopy Sciences). Sections were positioned between two ACLAR sheets (Electron Microscopy Sciences) and cured at 55 °C for 72 hours.

The ROI, CA1 stratum radiatum, was excised from the flat-embedded sections on ACLAR sheets and affixed to the tops of resin blocks. Ultrathin sections (∼75 nm) were cut using an ultramicrotome (ARTOS 3D, Leica Biosystems) and collected on silicon nitride chips. Sections were imaged using a Crossbeam 350 field emission scanning electron microscope with a Gemini column (Zeiss), operated at 1.4 kV and 1.2 nA. Microglial cell bodies were identified by their overall shape, characteristic heterochromatin pattern pockets, and long narrow stretches of endoplasmic reticulum ^42^. Immunohistochemical labelling was identified on microglial cells as electron-dense puncta localized to endosomes—spherical structures containing numerous intraluminal vesicles (ILVs), often associated with lysosomes (spherical, single membrane-bound, electron-dense organelles with heterogeneous granular luminal contents). Labelled microglia were imaged at 5 nm resolution, and images were exported in TIFF format using the Zeiss ATLAS Engine 5 software (Fibics).

### CLOZAPINE TREATMENT AND PHARMACOLOGICAL ANALYSIS

#### Clozapine administration via diet

To test the effects of antipsychotic treatment in our mouse model of acute psychosis, clozapine was chronically administered via specially formulated chow using a gradual dose titration. This approach mimics clinical dosing protocols to minimize sedation and other adverse effects. Diets were commercially prepared at increasing concentrations of 125, 250, 375, 500, 625, and 750 ppm of clozapine in standard rodent chow; control animals received identical base diet without clozapine. The titration period lasted ten days, with mice advancing to the next higher concentration every two days. On each titration day, mice were weighed before the food hopper was emptied and refilled with the next clozapine diet, allowing close monitoring of body-weight changes during dose escalation. Following completion of the titration period, mice were maintained on 750 ppm chow for two days, before subcutaneous mRNA-LNP immunizations were initiated according to the schedule described above. Higher doses were not pursued due to mortality observed in approximately 10% of animals without any prior or post-mortem signs of ill health, consistent with sudden cardiac death arising from the known pro-arrhythmic effects of antipsychotic drugs.

#### Plasma clozapine concentration measurement with LC-MS/MS

To assess clozapine exposure, we measured plasma clozapine concentrations by LC-MS/MS using an adapted method.^43,44^ Mouse plasma (15 µL, when available or 7.5 µL) was extracted with 60 µL or 30 µL ice-cold methanol:acetonitrile (16:84, v/v) containing a serial dilution of clozapine-d8 internal standard (3.9-125 ng/mL; ChemCruz), then vortexed and centrifuged at maximum speed for 10 min at 4 °C; supernatants were transferred to LC-MS vials with glass inserts and immediately analyzed, with pooled biological QC samples and calibration standards included in each batch. Extracts (5 µL) were injected onto a HyPURITY C18 column (50 × 2.1 mm, 3 µm; Thermo Scientific; Cat# 22103-052130) mounted on a Dionex UltiMate 3000 LC system and separated using a 1.5-min gradient from 0-100% solvent B, followed by a 0.7 min wash and 0.4 min re-equilibration at 500 µL/min (column 20 °C; autosampler 4 °C). Solvent A was water containing ammonium acetate (250 mg/L), acetic acid (1.75 mL/L), and trifluoroacetic anhydride (100 µL/L), and solvent B was acetonitrile containing the same additives. Detection was performed on a TSQ Quantiva (Thermo Scientific) in positive ESI mode (spray voltage 3.5 kV; probe 350 °C; sheath gas 50 a.u.; auxiliary gas 15 a.u.; collision gas 1.5 mTorr; Q1/Q3 resolution 0.7/1.2 FWHM) using SRM transitions m/z 327 → 270 (clozapine) and m/z 355 → 275 (clozapine d8); data were acquired in Xcalibur (v4.0.27.10; RRID:SCR_014593) and quantified in TraceFinder (v5.1) using internal standard calibration.

#### ELISA for anti-nuclear autoantibodies

To detect IgG antibodies against DNA and RNA in our mouse model of lupus, an indirect enzyme-linked immunosorbent assay (ELISA) was employed. MaxiSorp 96-well plates were pre-coated with Poly-L-Lysine (0.1% w/v, 1:50 in DEPC-treated water; 100 µL/well) and incubated for 4-5 h at room temperature in a humid chamber, then washed by flicking and blotting. Plates were coated overnight at 4 °C with 2.5 µg DNA (Sigma-Aldrich, D7290) or yeast RNA (Ambion, AM7120G) diluted in coating buffer (1% BSA in PBS, 50 µL/well), with buffer alone in control wells. After wash with PBS-T, wells were blocked with 150 µL/well blocking buffer for 2 h at room temperature and washed three times with PBS-T. Serum samples diluted in blocking buffer (diluted 1:40, at 50 µL/well) were added and incubated overnight at 4 °C, followed by five washes with PBS-T. Goat anti-mouse IgG-alkaline phosphatase (1:2,000 in blocking buffer; 50 µL/well) was added and incubated for 1 h at 37 °C, and plates were then washed five times. Colorimetric signal was developed by adding 50 µL/well alkaline phosphatase substrate (phosphatase tablets, 1 mg/mL in developing buffer) and incubating for 10 min at 37 °C, and optical density was measured at 405 nm with a 605 nm reference using a microplate reader.

### QUANTIFICATION AND STATISTICAL ANALYSIS

#### General statistical approach

All statistical analyses were performed in MATLAB 2024b (MathWorks). Statistical significance was defined as p < 0.05 (two-sided) throughout. For two-group comparisons, Welch’s t-tests were applied to account for unequal variance in the groups, except for antibody concentration data, where non-parametric Wilcoxon rank-sum tests were used to account for floor effects at the assay detection limit. For paired comparisons of the usage proportion of each behavioral syllable measured across two groups (Fig. 1Q and 3H), a paired t-test was used. For repeated-measures designs in which multiple observations were nested within animals, linear mixed-effects models (LMEs) were used to account for within-subject correlations; mouse was included as a random intercept in all LME models, and statistical significance was assessed by ANOVA of the fitted model, followed by post-hoc Welch’s t-tests.

To minimize animal use in accordance with the 3Rs, experiments presented in Figs. 1 and 6 were prospectively designed as independent comparisons sharing a common reference group control. These experiments were run simultaneously as a single combined cohort with NMDAR immunized mice as a shared reference group compared against control immunized mice in Fig. 1 and against clozapine-treated NMDAR immunized mice in Fig. 6. Moreover, in Figure 2 wild-type NMDAR immunized mice showed an unusually high rate of dropout due to reaching humane endpoints (39% compared to 8% in other experiments). To obtain more precise estimates of home cage activity in this group, this cohort was supplemented with NMDAR-immunized animals from Figures 1 and 6, collected under otherwise identical conditions.

#### Home cage locomotion data analysis

Home cage locomotor activity was continuously recorded throughout the experimental period using the Tecniplast DVC interface and analyzed to assess the effects of immunization on spontaneous activity.

To remove changes in home cage activity unrelated to the experimental manipulation, several periods were excluded from analysis. Data prior to three days before the first immunization were discarded as this represented the habituation period to the Tecniplast DVC cages. A transient decrease in activity was noted following each injection in both the control and the NMDAR group, therefore data from this four day period was discarded. DVC Tecniplast racks do not produce reliable activity data from electrode 2, so this electrode was not included in locomotion analysis.

Locomotion was assessed using DVC cages, which estimate mouse position via capacitance measurements every 0.25 seconds; distance travelled per minute was used as the measure of locomotion. Analysis was restricted to the dark phase (19:00–07:00), when mice are most active and were undisturbed by experimental interventions. A locomotion index was calculated by normalizing each animal’s daily distance to its mean over the three days preceding the first immunization, to account for individual differences in baseline activity. For the correlation between CSF antibody levels and locomotion presented in Figure 4C, mice were placed in DVC cages one week after the first immunization; raw distance travelled per minute was therefore presented.

For time course plots, data were averaged weekly per animal before calculating group means. For summary plots, data were averaged across the post-immunization window, defined as day 29 until the end of the experiment, excluding the four-day period with transient decrease in activity following each injection (see above). For Figs. 1, 2 and 6, the post-injection exclusion period spanned day 29-32. For Fig. 5, the final immunization was delayed, shifting the post-injection exclusion period to days 36-39; the post-immunization window was otherwise defined equivalently.

#### Motion sequencing (MoSeq) data analysis

To characterize the effects of immunization on the structure of spontaneous behavior, Motion sequencing (MoSeq) data were acquired with a depth camera and processed with the MoSeq software (https://github.com/dattalab/moseq2-app).

To assess open field activity, mouse centroid position on each frame was extracted, and the following measures were calculated: **Distance traveled**: To assess locomotion in the open field, trajectory length was calculated as the sum of the length of trajectory segments. **Stereotypic cycling**: To quantify stereotypical circling, we defined the ‘loop area’ as the area enclosed when the mouse crossed its previous path, restarting the measurement after each crossing. Area was calculated using the shoelace formula, and the mean of loop areas per session was calculated.

Open field measures from the final post-immunization timepoint at week 7 are presented for the mRNA immunization experiments (Figures 1 and 6). For the passive transfer experiment (Figure 3), two mice (one per group) were excluded due to poor postoperative recovery, evidenced by progressively decreasing distance traveled throughout the experiment. Remaining mice were analyzed with a linear mixed effects model for each metric (distance and loop area) fitted to all three sessions with formula:

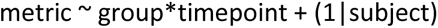

The effect of group was significant so differences were tested at each timepoint using Welch’s t-test.

To assess the behavioral repertoire, behavioral sequences were parsed into syllables by fitting autoregressive models to the data. As per the procedure described previously^37^ a range of models were fitted with different values of kappa (which determines the syllable length) and the kappa value with modal syllable length most closely matching the model-free estimate of syllable changepoints was chosen. 100 autoregressive models with this kappa value were fitted and the one with highest median log likelihood was selected. ‘Crowd movies’, showing 20 instances of mice performing each syllable, were visually inspected and each syllable was labelled with a verbal category by one author (HRF). Five out of 100 syllables were excluded from the model fitted to data from the mRNA immunization experiments because a crowd movie was not produced, or the syllable showed high variability of behaviors.

For the mRNA immunization experiment, a single model was fitted to all data from all timepoints, and all post-immunization timepoints were analyzed statistically to increase power when fitting a model with multiple factors and levels. Mean syllable usage across all post-immunization sessions is displayed. For passive transfer experiments, all sessions were fitted and analyzed statistically. Behavioral syllables were analyzed to derive the following measures:

##### Syllable usage

For each session, the proportion of time spent in each behavioral syllable was calculated. Since these usage proportions across syllables necessarily sum to one, changes in one syllable mathematically constrain the other thereby violating the assumptions of standard statistical tests. To account for this compositional nature of the data, the data were centered log-ratio transformed before statistical analysis. For Figure 6, clozapine-treated compared to untreated mice showed baseline differences in behavioral composition making absolute values difficult to interpret; data were therefore normalized to the group mean of the pre-immunization session. This rendered the data non-compositional so center log ratio transformation was not performed.

A linear mixed effects model was fitted with formula:

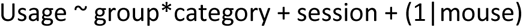

ANOVA of this model was performed. Comparison between groups for each category were performed with Welch’s t-tests.

##### Syllable velocity

To enable direct comparison with previous work,^5,16^ we analyzed the relationship between syllable velocity and usage. The mean velocity of each syllable and mean usage proportion of each syllable were calculated separately for the experimental groups. Syllables were ranked by mean velocity. For Figures 1P and 3G, the ratio of usage proportion in the NMDAR-immunized group to the control group was log transformed and plotted against velocity rank. For Figures 1O and 3H, the usage proportion of the 15 highest velocity syllables was compared between groups using a paired t-test.

##### Entropy

To analyze the entropy of behavioral sequences, a transition matrix was derived for each session by calculating the probability of transitioning from each syllable to every other syllable on consecutive frames. Syllable self-transitions were set to zero, and Shannon entropy calculated using MATLAB *entropy* function. Entropy was exclusively calculated for sessions where objects were present in the arena, specifically, sessions where the novel object recognition test was conducted in the MoSeq setup. This approach was adopted following previous work showing group differences in entropy only when objects were present in the arena.^46^ No baseline data with objects in the arena was available. Mean entropy from the familiarization and testing sessions was calculated for each mouse and groups were compared with Welch’s t-test.

#### Novel object recognition data analysis

We calculated the novel object recognition index to quantify the preference for novel objects over familiar objects. In line with previous work,^47^ object proximity was defined as any instance in which a body part was within a 10 cm radius of the center of an object. Object exploration was defined as the orientation of the nose toward the object during object proximity. The novel object recognition index was calculated as the difference between the time spent exploring the novel object and the time spent exploring the familiar object, divided by the total exploration time.

For the mRNA immunization experiment, data was available from a single timepoint at week 7 and groups were compared with Welch’s t-test.

For the passive transfer experiment, data were available from days 4, 9 and 13 post implantation. An LME model was fitted with formula:

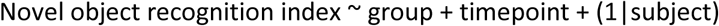

ANOVA of this model was performed. Mean data per mouse across all timepoints is plotted.

#### Prepulse inhibition data analysis

To quantify prepulse inhibition of the acoustic startle response, raw voltage recordings from the SR-LAB piezoelectric sensor were exported into MATLAB. Voltage recordings from 50ms to 150ms post-stimulus were analyzed to capture the peak startle response to the startle stimulus. Maximum voltage in this period was extracted and the prepulse inhibition was calculated as:

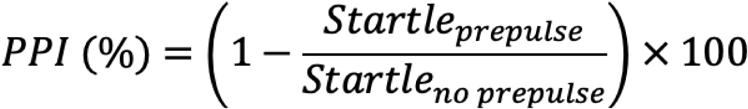

Prepulse inhibition was averaged within each session across all startle stimulus levels. Groups were compared with Welch’s t-test.

#### Correlations between behavioral metrics and autoantibody levels

Pearson’s correlation coefficient was used throughout. To identify the timepoint at which terminal autoantibody levels were most predictive of home cage behavior, the serum autoantibody concentrations measured at the experimental endpoint were calculated with daily home cage activity values across the full experimental timecourse.^48^ This revealed that correlation was maximal around day 37 for all home cage metrics; therefore data from this timepoint are presented.

To assess the relationship between autoantibody levels and behavior more broadly, 13 correlations were calculated between serum autoantibody concentrations and all behavioral metrics shown in Figure 1, using Pearson’s correlation coefficient to test for significance.

#### Anti-NMDAR antibody ELISA data analysis

A surrogate standard curve, generated using a purified monoclonal anti-NMDAR antibody of known concentration, was run on every ELISA plate alongside experimental samples and fitted with a 4-parameter logistic (4PL) curve. The range between the upper and lower bounds of the logistic curve was calculated. Lower limit of quantification was set at 5% of the range above the lower limit and upper limit of quantification was set at 5% below the upper limit. Data values that fell outside of the quantification range were censored at the lower or upper concentration limit and retained for group-level analysis rather than excluded, to avoid systematic bias in group-level estimates given the small sample size. Antibody concentrations are expressed as equivalent concentrations relative to the reference monoclonal antibody. Non-parametric Wilcoxon rank-sum tests were used for all group comparisons of antibody concentrations, as a substantial proportion of control group concentration fell, as expected, at or below the lower limit of quantification, introducing floor effects.

#### Flow cytometry data analysis

Flow cytometry data were acquired using FlowJo software and exported for downstream analysis.

Immune cell populations were identified by sequential gating on live (Zombie-NIR⁻), singlet excluded the circling immune cell in the blood, CD45 PE⁺ cells. Microglia were defined as CD45-Low, CD11b^+^. Macrophages were defined as CD45-Hi, F4/80⁺CD64⁺. From the CD45-Hi, F4/80⁻CD64⁻ (non-macrophage) fraction, dendritic cells were identified as MHC-II⁺CD11c⁺, and NK cells and neutrophils were separated as NK1.1⁺Ly6G⁻ and NK1.1⁻Ly6G⁺, respectively. Among the remaining CD45-Hi cells, B cells were defined as CD19⁺CD3⁻, while T cells were gated as CD19⁻CD3⁺ and further subdivided into CD4⁺ T cells (CD4⁺CD8⁻), CD8⁺ T cells (CD4⁻CD8⁺), and other T cells (CD4⁻CD8⁻).

Absolute cell counts in brain were calculated using the counting beads method (51,000 beads added per sample) according to the formula: absolute count = (population count / bead count) × 51,000. Five categories of measurements were analyzed: (1) raw flow counts for each population, (2) percentage of parent population, (3) geometric mean fluorescence intensity for specific markers, (4) absolute cell counts, and (5) percentage relative to total CD45-Hi cells. Welch’s t-tests were used for all group comparisons.

#### FFPE brain tissue immunofluorescence microscopy data

Images were analyzed in QuPath (v0.5.1; RRID:SCR_018257). The hippocampus was manually annotated, and mean fluorescence intensity was quantified within the hippocampal ROI for the IgG and NMDAR channels. Welch’s t-tests were used for all group comparisons.

#### Floating brain tissue immunofluorescence microscopy data

3D reconstruction and analysis of Iba1-positive microglia were performed in Imaris 10.2 (Oxford Instruments). Machine learning segmentation was carried out using a manually trained classifier in which three representative cells per slide were annotated as foreground and background. Segmentation outputs were inspected for each slide, and cells with incomplete somata, cells fused to neighbouring microglia, and cells truncated at image borders were excluded. Surface reconstructions were then used as templates for 3D filament tracing. Filament detection parameters were adapted from published microglial morphometry workflows^49^ and applied using the Imaris Filament tool, with starting-point detection set to a largest diameter of 3.95 µm and seed points of 0.3 µm, and seed-point removal set to a sphere diameter of 15 µm. Filament data for each microglial cell were exported to MATLAB. Linear mixed effects models were fitted to the length and area data from all cells with fixed factor group and random factor mouse, and an ANOVA was performed. Mean value per mouse is plotted

#### Western blot data analysis

Images were analyzed using ImageStudio v 6.1.0.79 (LiCor Biotech). Chemiluminscent images were inverted before further analysis. Rectangular regions-of-interest (ROIs) were drawn around bands of interest for quantification. For total protein staining, rectangular ROIs were drawn across the whole length of the lane. Background was defined as the median of three pixel wide area above and below the ROI of interest. Signal was calculated as the sum of foreground pixel intensities minus the ROI area multiplied by the median background intensity. Band signals were normalized to the lane-specific total protein signal. Two lanes with total protein level <50% of the maximum (indicative of loading failure) were excluded from the analysis.

A linear mixed-effects model was fitted to the data with group as a fixed effect and mouse as a random effect to account for the two brain hemispheres from each mouse. Statistical significance was assessed by ANOVA of this model. Mean signal per mouse is plotted. For presentation purposes, image brightness and contrast were uniformly adjusted, and lanes were reordered in Adobe Illustrator to group cases and controls separately.

#### Clozapine level analyses

Mice with a serum clozapine level <15ng/ml at the end of the experiment were removed from the analysis. Three mice from the behavioral cohort and two mice from the immunophenotyping cohort were excluded.

For antibody concentration statistics in the clozapine experiments, where all mice were given anti-NMDAR immunizations, concentrations below the lower limit of quantification were excluded.

#### ELISA for anti-nuclear autoantibodies data analysis

Autoantibody levels are reported as raw optical density (OD) values. Group comparisons were performed using Welch’s t-test.

