## Supplementary Videos for "The antipsychotic drug clozapine suppresses autoimmunity driving psychosis-like behavior in mice"

### Slide 1
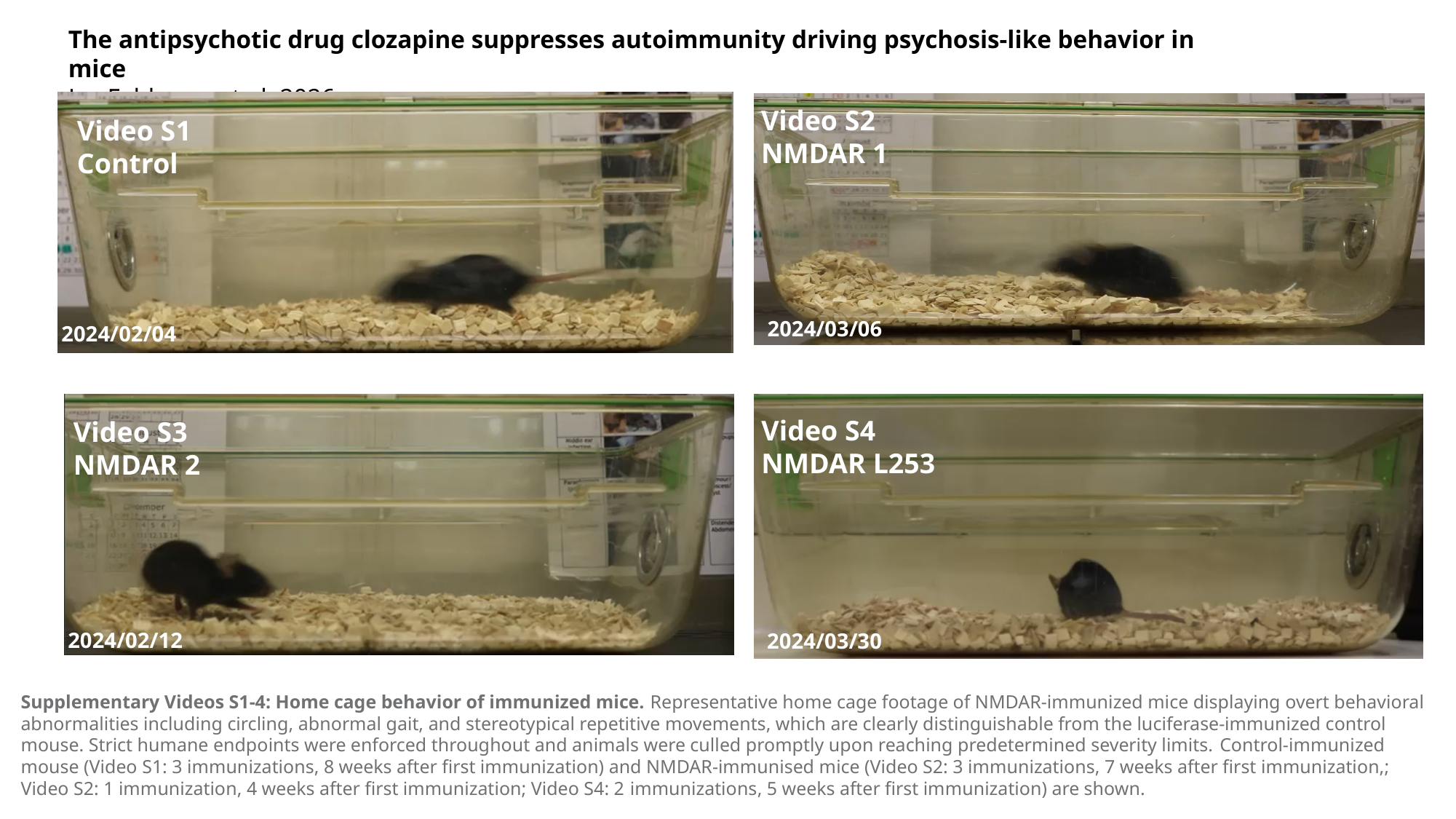

#### The antipsychotic drug clozapine suppresses autoimmunity driving psychosis-like behavior in mice Le, Feldman, et al. 2026
Video S2
NMDAR 1
Video S1
Control
NMDAR L225
2024/03/06
2024/02/04
Video S4
NMDAR L253
Video S3
NMDAR 2
2024/02/12
2024/03/30
2024/02/12
Supplementary Videos S1-4: Home cage behavior of immunized mice. Representative home cage footage of NMDAR-immunized mice displaying overt behavioral abnormalities including circling, abnormal gait, and stereotypical repetitive movements, which are clearly distinguishable from the luciferase-immunized control mouse. Strict humane endpoints were enforced throughout and animals were culled promptly upon reaching predetermined severity limits. Control-immunized mouse (Video S1: 3 immunizations, 8 weeks after first immunization) and NMDAR-immunised mice (Video S2: 3 immunizations, 7 weeks after first immunization,; Video S2: 1 immunization, 4 weeks after first immunization; Video S4: 2 immunizations, 5 weeks after first immunization) are shown.
